# Cancer-induced Nerve Injury Unveils a Sympathetic-to-Sensory Nerve Axis in Head and Neck Cancer

**DOI:** 10.1101/2025.09.19.677339

**Authors:** Andre A. Martel-Matos, Lisa A. McIlvried, Nicole L. Horan, Nicole A. Rodriguez, Jared I. Rothberg, Megan A. Atherton, Stephen V. Glass, Marci L. Nilsen, Nicole N. Scheff

## Abstract

Oral squamous cell carcinoma (OSCC) is one of the most painful cancers, with patients frequently reporting spontaneous, neuropathic-like pain. While sympathetic and sensory nerves have been individually implicated in cancer progression, whether and how these systems interact to drive pain and tumor growth has remained unclear. Here, we integrate prospective human data with reverse-translational mouse models to reveal that cancer-induced nerve injury unveils crosstalk between sympathetic postganglionic neurons and trigeminal sensory afferents in the tumor microenvironment. In patients, circulating norepinephrine (NE) correlated with spontaneous pain and perineural invasion, identifying a potential sympathetic contribution to disease burden. In mice, aggressive non-immunogenic OSCC tumors evoked spontaneous nociceptive behaviors, elevated tumoral NE, and sensory nerve injury marked by ATF3 expression and hyperexcitability. Tumor-associated sensory neurons acquired adrenergic sensitivity through α1-adrenergic receptor plasticity, while sympathetic neurons exhibited plasticity characterized by sprouting, altered gene expression, and heightened excitability, creating a maladaptive feed-forward loop that amplified nociceptive signaling. Disrupting this sympathetic-sensory communication by sympathectomy or selective ablation of TRPV1⁺ sensory fibers reduced tumor growth, sympathetic tone, and spontaneous pain like behaviors, although sensory adrenergic sensitivity persisted. Together, these findings establish that reciprocal sympathetic-sensory plasticity and crosstalk in the tumor may fuel both OSCC progression and neuropathic like pain. Targeting this peripheral neuroplasticity may offer a translational strategy to limit tumor growth and alleviate pain.

**One Sentence Summary:** Reciprocal plasticity and crosstalk between sympathetic and sensory nerves drive both tumor progression and neuropathic-like pain in oral squamous cell carcinoma, suggesting peripheral neuroplasticity as a therapeutic target.

## 1.0 Introduction

The cancer neuroscience field has explored the role of the peripheral nervous systems in the context of tumor growth and anti-tumor immune response, albeit largely focused on autonomic signaling. Sympathetic nervous system (SNS) signaling, facilitated by the release of norepinephrine (NE), has been found to play a direct role in promoting tumor growth, cell survival and metastasis through beta adrenergic receptor expression on tumor cells (*1, 2*). Moreover, sympathetic activation can also modulate immune responses and potentially contribute to immune evasion by the tumor (*3–5*). Recently sensory nerve activity, with a particular focus on TRPV1-expressing nociceptors, has also been implicated in promoting tumor immunosuppression, at least in part through the release of neuropeptides such as calcitonin gene-related peptide (CGRP) (*6, 7*).

Although the peripheral nervous systems have mostly been studied independently, emerging evidence suggests that these systems may interact in the context of cancer to jointly influence tumor progression. In a mouse model of breast cancer, both pharmacogenetic and chemogenic activation of the central medial amygdala (CeM) led to increased anxiety-like behaviors and accelerated tumor growth (*8*). While critical for emotional processing and fear regulation, the CeM also serves as a key hub for nociceptive (i.e. pain) processing from both trigeminal and spinal nerves. Importantly, activation of sensory nerves by the tumor microenvironment (TME) may precede the onset of autonomic nervous system responses; transneuronal tracing studies in an orthotopic oral squamous cell carcinoma (OSCC) model found that tumor-associated trigeminal neurons activate central circuits involved in both pain and stress processing, and analgesic use can attenuate anhedonia behavior (*9*). Together this suggests that the activation of sensory nerves by the TME may send ascending signals to the central nervous system (CNS), which in turn could modulate descending autonomic signaling to the TME. Despite evidence of CNS communication, peripheral sympathetic-to-sensory nerve axis within the TME remains unexplored.

There is evidence of both sensory and sympathetic activation during cancer. Many cancer types result in pain due to activation of sensory nociceptive afferents in the TME. OSCC is considered one of the most painful cancer types with 80% of patients reporting pain at diagnosis (*10, 11*). Furthermore, OSCC tumor growth is often aggressive, with advanced disease defined by destruction of surrounding tissue (*12*) (i.e. deep muscle, medullary bone (*13*) as well as perineural invasion (PNI) (*14*). Cancer-associated pain is considered mixed pathophysiology, including nociceptive and neuropathic (nerve injury) pain components (*15*). Neuropathic cancer pain, with a prevalence of 54% in patients with locally advanced oral cavity tumors (*16*), is due to either tumor-induced nerve compression or infiltration, or is secondary due to changes in the perineural environment in response to tumor-secreted cytokine dysregulation (*17, 18*). Additionally, systemic sympathetic hyperactivity has been noted in patients with HNSCC compared to non-cancer patients (*19*), and stellate ganglion blocks have been used in HNSCC patients to successfully attenuate cancer-related facial pain (*20*).

Clinical and preclinical neuropathic pain studies suggest direct and indirect peripheral crosstalk between sensory and sympathetic fibers at sites of neural insult. Xenograft and syngeneic OSCC mouse models revealed that tongue tumor growth can drive a robust nerve injury response in tumor-associated sensory neurons (*21, 22*). Following injury, sensory neurons have been shown to develop heightened sensitivity to catecholamines (*23*), indicating a direct mechanism for sympathetic-sensory crosstalk. An indirect mechanism has been implicated as well; non-selective beta-adrenergic receptor antagonism reduced orofacial nociceptive behaviors via tumor necrosis alpha (TNFα) release from tumor cells (*1*). Together, these findings suggest a likely involvement of SNS signaling impacting not only tumor progression but also the maintenance of tumor-associated sensory neurons and subsequent neuropathic pain processing.

We hypothesize that sympathetic signaling secondary to oral cancer-induced nerve injury drives neuropathic pain via adrenergic sensitivity in tumor-associated sensory neurons. In the present study, we use a prospective observational study and reverse translate in mouse models to investigate the mechanistic keystones of sensory and sympathetic nerve communication on tumor progression and associated pain. Our findings show that cancer-induced nerve injury results in sympathetic-sensory neuronal plasticity and crosstalk in the TME that exacerbates tumor progression and oral cancer-associated neuropathic pain. Understanding peripheral nerve plasticity and communication during tumor progression will aid in identifying patients prior to treatment who will benefit most from adjuvant therapeutic strategies that modulate the peripheral nervous system to improve outcomes.

## 2.0 Results

### 2.1 Elevated Circulating Norepinephrine (NE) in Oral SCC Patients Correlates with Spontaneous Pain

Previous studies have indicated that circulating norepinephrine (NE) is elevated in head and neck cancer patients compared to controls, suggesting sympathetic dysregulation (*19*). Comorbidities such as pain, depression, and anxiety are commonly reported by cancer patients (*24*) and are associated with increased sensory and sympathetic neurotransmission activity (*25, 26*). To begin to examine the potential role of sympathetic signaling in oral cancer pain, we used a prospective observational study executed to test the hypothesis that cancer-associated pain and psychological symptom burden correlate with local sympathetic nerve presence and plasma catecholamine levels in HNSCC patients. The majority of patients eligible for final analysis had SCC tumors in the oral cavity (*n* = 88), with 8 additional patients with tumor presentation in the oropharyngeal and neck regions. The majority were male (*n* = 60, 62.5%) and white (*n* = 92, 96%) with a mean ± SD age of 67.5 ± 13.0 years. Primary tumor stage was identified by a clinical pathologist as I/II (n=58, 60.4%) and nodal stage of 0/1 (n=71, 74.0%). Additional demographics and clinical characteristics are shown in **Table 1**. General head and neck pain scores were collected using the Brief Pain Inventory (BPI), which provides a summation score for pain severity and pain interference (*27*). Specific cancer-related pain scores were assessed using the University of California Oral Cancer Pain Questionnaire (OCPQ), which provides a composite score as well as function-evoked pain (i.e. while talking and eating) and spontaneous/going pain (i.e. while at rest) (*28*). There was an increase in patient-reported general and oral cancer-specific pain with primary tumor stage (**Fig 1A,B**). Patient-reported psychological symptoms were also assessed using the Patient Health Questionnaire (PHQ-8) and General Anxiety Disorder 7 questionnaire (GAD-7); there was no difference in reported depression and anxiety scores between primary tumor stage (**Fig 1C**). First, we explored the relationship between patient-reported pain and psychological symptoms in HNSCC patients prior to surgery. Using univariable analysis, there was a positive correlation between anxiety and OCPQ composite score (p=0.017, 95% CI [0.131, 1.180]); however, this effect was lost when controlling for age, sex, and tumor stage. Instead, patients who experience higher levels of mean pain severity (i.e., BPI) or oral cancer pain (i.e., OCPQ composite score) were more likely to experience higher levels of depression. On multivariable analysis, when controlling for age, sex, and stage, the positive association between depression and both OCPQ composite score and BPI mean pain severity remained (p=0.004, p=0.048, respectively).

**Figure 1:**
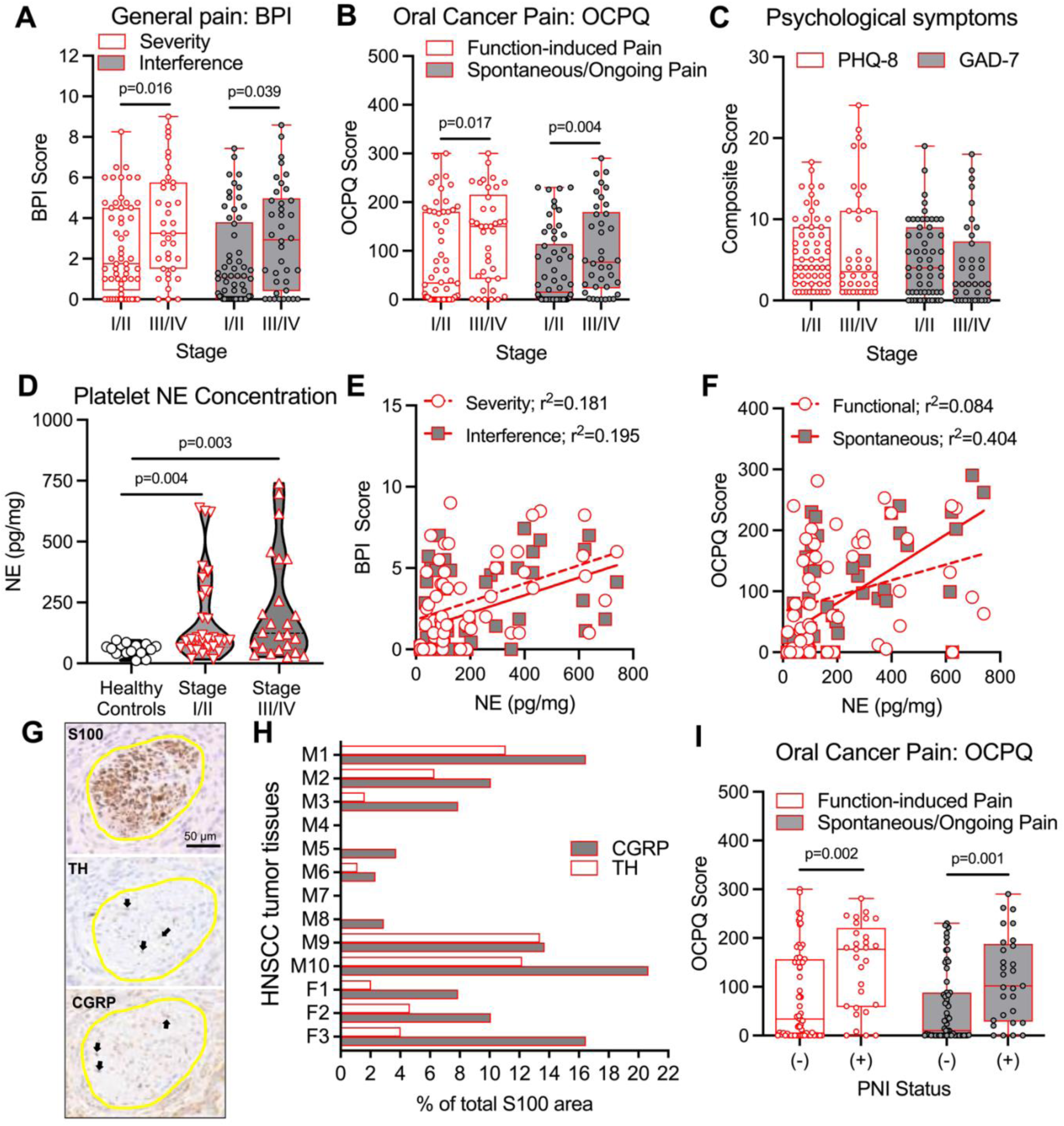
Elevated circulating norepinephrine in OSCC patients correlates with spontaneous pain. (A-C) Pain and psychological symptom burden as measured by the symptom-specific questionnaire tools in 96 HNSCC patients. Statistical significance was measured using Mann-Whitney U test within symptom subtype. P values are denoted on the graph. (A) Generalized pain was measured by the Brief Pain Inventory (BPI). Pain severity (white bars) and pain interference (gray bars) scores are represented as box plots with each data point representing a patient pain score. (B) Oral cancer-specific pain was measured by the OCPQ. Function-evoked (i.e. while using oral cavity) (white bars) and spontaneous/ongoing pain (gray bars) are represented as box plots with each data point representing a patient pain score. (C) Psychological symptom scores were measured as depression using the PHQ-9 tool (white bars) and anxiety using the GAD-7 tool (gray bars); represented as box plots, each data point representing a patient pain score. (D) Circulating norepinephrine was measured by ELISA in blood platelets isolated prior to treatment in 12 healthy controls and 56 HNSCC patients stratified by tumor stage. Kruskal-Wallis test with Dunn’s multiple comparisons test, P values are denoted on the graph. (E) The relationship between generalized pain severity (solid red line, white circles) and interference (dashed red line, gray boxes) as measured by the Brief Pain Inventory and circulating NE visualized by a simple linear regression in 56 HNSCC patients. (F) The relationship between function-evoked oral cancer pain (dashed red line, white circles) and spontaneous/ongoing pain (solid red line, gray boxes) as measured by the UCSF oral cancer pain questionnaire and circulating NE visualized by a simple linear regression in 56 HNSCC patients. (G) Representative image of S100 (top), tyrosine hydroxylase (TH, middle), and CGRP (bottom) positive immunoreactivity (IR) in HNSCC tumor section (Male, Stage IV). Nerve bundle is outlined in yellow. Positive stain is indicated by black arrows. Image magnification = 20x. (H) Quantification of the percent of total CGRP-IR and total TH-IR nerve area relative to total S100-IR nerve area across tumor tissue sections from 13 HNSCC patients. (I) Function-evoked (white boxes) and spontaneous/ongoing (gray boxes) oral cancer pain stratified by perineural invasion (PNI) pathology. Patients with PNI tumor pathology reported significantly higher pain compared to those without PNI. Mann-Whitney U test within pain subtype, P values are denoted on the graph.

**Table 1.** Demographic and Clinical Characteristics (n=96).

| **Variables** | **n(%); Median (Q1,Q3)** |
| --- | --- |
| **Age, years** | 70 (61, 76) |
| **Sex** |  |
| Male | 60 (63%) |
| Female | 36 (38%) |
| **Race** |  |
| White | 92 (96%) |
| Other | 4 (4%) |
| **Site** |  |
| Oral Cavity | 88 (92%) |
| Oropharynx | 6 (6%) |
| Tonsil | 1 (1%) |
| Neck | 1 (1%) |
| **Primary Tumor Stage** |  |
| TI-II | 58 (60%) |
| TIII-IV | 38 (40%) |
| **Nodal Status** |  |
| N0-1 | 71 (74%) |
| N2-3 | 25 (26%) |
| **Smoking** |  |
| No | 42 (44%) |
| Yes | 54 (56%) |
| **Alcohol** |  |
| No | 32 (33%) |
| Yes | 64 (67%) |
| **Perineural Invasion** |  |
| No | 61 (64%) |
| Yes | 29 (30%) |
| Not Evaluated | 6 (6.3%) |
| **Opioids** |  |
| No | 79 (82%) |
| Yes | 17 (18%) |
| **Brief Pain Inventory** |  |
| Severity | 2.3 (1.00, 4.8) |
| Interference | 1.4 (0.1, 4.4) |
| **Oral Cancer Pain (OCPQ)** |  |
| Composite | 193 (14, 431) |
| Function-evoked | 79 (5, 183) |
| Spontaneous | 41 (0, 143) |
| **PHQ-8** |  |
| Sum score | 4.0 (2.0,10.0) |
| **GAD-7** |  |
| Sum score | 3.0 (1.0, 8.0) |
| **Norepinephrine (NE)** |  |
| Concentration | 105 (67, 294) |
| Unknown | 39 |

Next, we sought to explore the relationship between cancer-associated pain and circulating NE concentration in HNSCC patients. Blood platelets were measured for NE by ELISA (n=58 cancer patients/n=15 healthy donor controls) as an overall measure of sympathetic signaling. Platelets were selected over plasma as a more robust metric for chronic sympathetic activity that is resistant to the acute confounding variables (*29, 30*). The median NE concentration was 103.9 (92.5, 163.9) pg/mg in HNSCC patients and 61.5 (43, 74.6) pg/mg in healthy controls (**Fig 1D**). Nonparametric analysis revealed an increase in platelet NE in HNSCC patients compared to healthy controls, irrespective of tumor stage (**Fig 1D**). When controlling for sex, age, primary tumor stage and perineural invasion (PNI), using a logistic regression model, we found that all pain scores were associated with NE concentration after adjusting for covariates, particularly spontaneous pain (**Fig 1E,F, Table S1)**; for example, a 1-point increase in OCPQ Spontaneous Score was associated with a 0.007 increase in natural log of NE concentration after adjusting for covariates (q < 0.001, 95% CI [0.004, 0.010]).

These findings indicate that circulating NE may contribute to patient-reported pain. While circulating levels of NE are not specific to the tumor microenvironment, they are likely representative of sympathetic postganglionic output in the periphery since the adrenal arm of the HPA axis primarily secretes epinephrine into circulation (*31*). To investigate this, we used PNI as a biological indication of potential sympathetic-sensory nerve communication; recent literature demonstrates that the majority of HNSCC tumor-associated nerves are mixed sympathetic-sensory nerve bundles (*32*). Consistent with these findings, nerve bundles identified retrospectively in patient tumor samples were predominantly composed of sensory (calcitonin gene-related peptide, CGRP⁺) and sympathetic (tyrosine hydroxylase, TH⁺) fibers; two of the 13 patients lacked TH-immunoreactivity, with only CGRP+ nerve bundles present in the tissue (**Fig. 1G,H, Suppl Fig 1A**). Within our prospective patient cohort, PNI, assessed by a clinical pathologist in resected tumor tissue, was detected in 29 (30%) patients; consistent with previous reports (*33*), patients with PNI pathology reported higher oral cancer pain compared to those without PNI (**Fig. 1I**). We explored the relationship between PNI and patient-reported pain and clinical variables. Univariable analyses showed that NE concentration and N-stage II/III were associated with odds of PNI (**Table S2**). While the odds of having PNI are 65% higher for HNSCC with stage III/IV compared to stage I/II, primary tumor stage was not significant when adjusting for multiple comparisons, likely due to the limited sample size. To assess what type of pain (i.e. function, spontaneous) serves as a predictor of PNI, the logistic regression model was used controlling for univariate-defined variables as well as sex and t-stage. The OCPQ composite score (OR: 1.0 95% CI [1,1.01], p=0.027) and spontaneous pain score (OR: 1.01 95% CI [1,1.02], p=0.011) were associated with PNI (**Table S3**). All together these data suggest a potential relationship between patient-reported pain, sympathetic signaling and tumor-associated peripheral nerves such that further exploration is warranted.

### 2.2 Syngeneic Orthotopic Mouse Oral Cancer Model Recapitulates HNSCC Patient Phenotype

To investigate the potential interaction between sympathetic and sensory systems in the context of oral cancer, we employed a reverse translational approach to recapitulate OSCC patient outcomes utilizing syngeneic orthotopic mouse models. Two different mouse oral cancer cell lines (MOC1 or MOC2) were injected in the anterior tongue of male and female C57BL/6 mice to produce tumors. These cell lines were selected to mimic two different phenotypes of OSCC based on tumor cell immunogenicity (*21, 34*) and indolent or aggressive tumor growth kinetics (*35, 36*); MOC1 elicits a robust tumor-associated immune response, with slower tumor growth and a maximum tumor burden around post inoculation day (PID) 40, while MOC2 induces a smaller immune response, with much faster tumor growth that peaks around PID14 (*21*) (**Fig 2A,B**). Both MOC1 and MOC2 tumor bearing mice experienced significant weight loss during tumor progression compared to sham mice (**Fig 2C**), suggesting decreased use of the oral cavity; meal analysis in tongue tumor mice has been utilized previously as an indication of nociception (*37*). Given that OSCC patients reported both function-evoked and spontaneous pain based on the OCPQ and BPI questionnaires, we sought to evaluate nociceptive behavior in the mouse tongue cancer models during orofacial activity (i.e. function-evoked pain) as well as at rest (i.e. ongoing, spontaneous pain). To evaluate nociceptive behavior associated with oral cavity activity, we utilized the dolognawmeter device and assay(*38*) to measure evoked nociceptive behavior through a discrete, operant gnawing task; an increase in gnaw-time is considered an index of pain and reversible by analgesics (*38*). Despite stark differences in growth kinetics, tumor growth significantly impacted gnawing behavior in both tumor models; MOC1 and MOC2 tumor mice exhibited over 400% increase in gnaw-time from baseline by PID40 and PID12, respectively, compared to time-matched sham controls (**Fig 2D,E**). To assess spontaneous pain in the absence of oral cavity activity, we utilized the Mouse Grimace Scale (MGS) (*39*) in tandem with the automated PainFace analysis platform (*40*) to quantify changes in facial features associated with grimacing behavior while minimizing bias. Following individual video recording sessions of sham, MOC1 and MOC2 tumor bearing mice, the first 200 usable video frames (defined as frames with ≥3 out of 4 facial action units detected: eyes, nose, ears, whiskers). Tongue tumor growth did not physically distort mouse facial features (**Supp 1B-D**). Facial action units were analyzed and visualized as a heatmap to represent changes in grimacing behavior over time. Grimacing behavior was measured using three methods: (1) area under the curve (AUC) analysis extracted from the heatmap, (2) average action unit score over 200 frames, and (3) the number of grimacing frames defined as the facial action unit score ≥1.5 standard deviations from baseline. Baseline recordings prior to tumor inoculation confirmed that all mice began with stable and low scores indicative of no grimace behavior (mean action unit score: 2.45; AUC=300). Unlike function-evoked nociceptive behaviors, only MOC2 tumor-bearing mice demonstrated spontaneous/ongoing nociceptive behaviors compared to sham mice with no differences between MOC1 tumor mice and sham (**Fig 2F,G, Supp 1E-G**); a full-time course of MGS nociceptive behaviors in MOC2 tumor bearing mice revealed a similar onset to tumor growth and evoked nociceptive behavior. We found no significant effect of sex in tumor-bearing mice at PID12 (**Supp 1E-H**).

**Figure 2:**
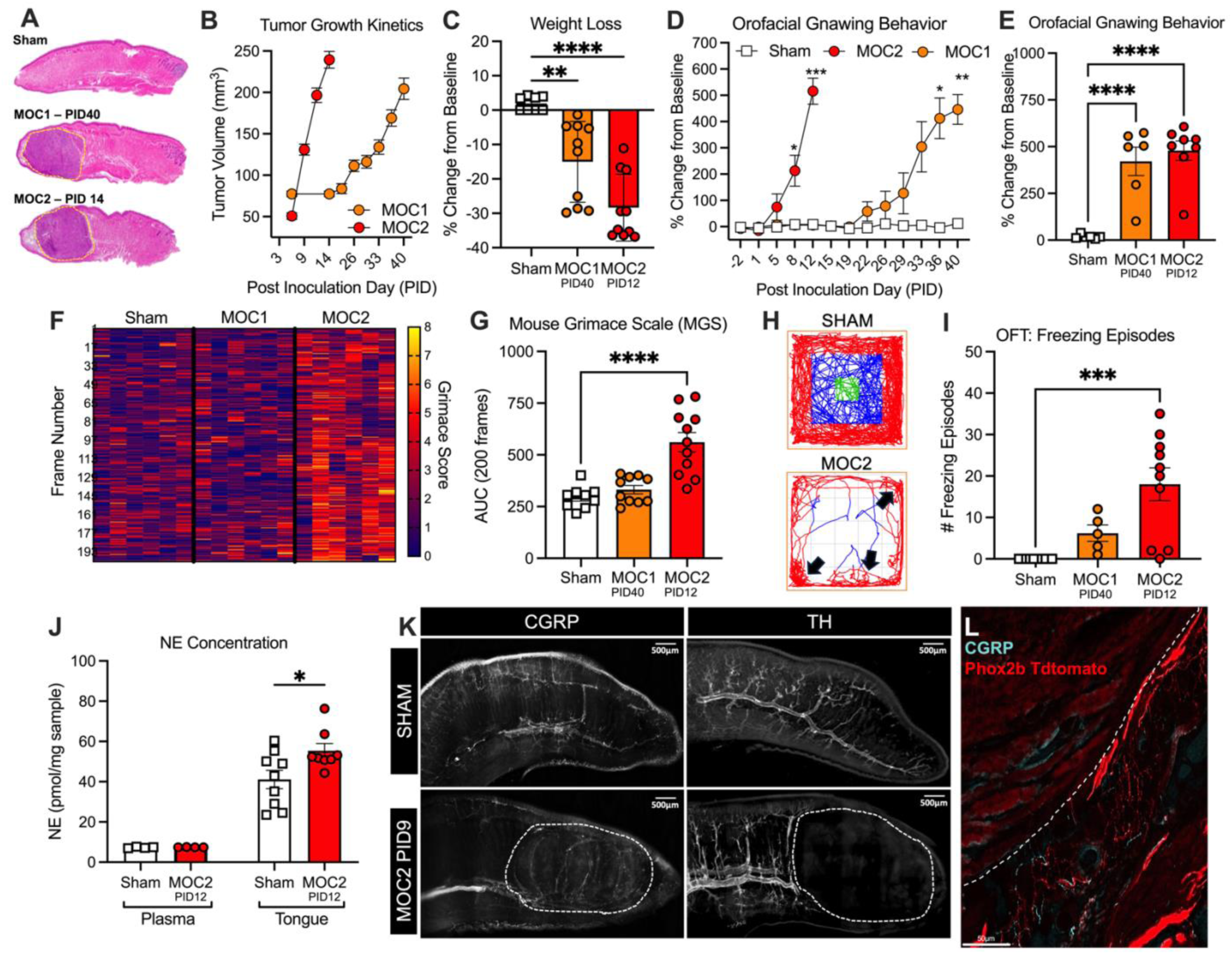
Syngeneic orthotopic mouse oral cancer model exhibits evoked and spontaneous pain behaviors. (A) Orthotopic tongue tumors were established via anterior injection of MOC1 (1×10⁶ cells) or MOC2 (2×10⁴ cells) cells into the tongues of mice. Representative hematoxylin and eosin images (5µm) for sham, MOC1 PID 40 and MOC2 PID12 tongues. Tumor bordered is indicated by yellow hashed line. PID = post inoculation day. (B) Tumor growth kinetics were established through caliper measurement in MOC1 (weekly) and MOC2 (every 3 days) tumor bearing mice starting at PID7. n=6/sex/group. (C) Tongue tumor-induced changes in body weight. Cancer weight loss was calculated as a percent change in body weight at a timepoint corresponding to equivalent tumor burden between MOC1 and MOC2 models, compared to baseline, pre-inoculation weight. n=6/sex/group, One-way ANOVA, **p<0.01, ****p<0.0001. (D-E) Operant gnawing behavior using dolognawmeter as a measure of evoked nociceptive behavior. Percent change from baseline gnaw-time in sham and MOC1 and MOC2 tumor-bearing mice plotted over time (C) and on peak pain days when tumor size was matched across models (D). n=3-4/sex/group, Two-way ANOVA, Cancer by time interaction (C), and One-Way ANOVA (D), *p<0.05, **p<0.01, ***p<0.005, ****p<0.0001. (F-G) Mouse grimace scale (MGS) as a measure of spontaneous nociceptive behavior. Data is visualized as a heatmap of individual grimace scores (range 0–8) from the first 200 usable frames per mouse (F). Quantification was calculated as the area under the curve (AUC) for grimace scores across the first 200 frames (G). n=4-5/sex/group, One-way ANOVA, ****p<0.0001. (H-I) Open Field Test (OFT) as a secondary measure of spontaneous nociceptive behavior. Representative track plots with freezing episodes indicated by black arrows (H). Analysis was done through AnyMaze software to quantify the total number of freezing episodes in which a mouse lacks any movement for ≥3 seconds during a 20-minute recording session. n=2-5/sex/group, One-way ANOVA, **p<0.01. (J) Norepinephrine concentration in plasma and tongue tissue from sham and MOC2 tumor bearing mice at PID12 quantified by mass spectrometry. n=4-5/sex/group, Independent T-test, *p<0.05. (K) Representative images of sensory and sympathetic nerve innervation in mouse tongue tissue at PID9. Sagittal tongue sections (300 µm) from sham and MOC2 tumor-bearing mice were optically cleared using the CUBIC protocol, immunostained for calcitonin gene-related peptide (CGRP; sensory fibers) or tyrosine hydroxylase (TH; sympathetic fibers), and imaged as stitched z-stacks under full focus. (L) Representative image of CGRP staining in a 20µm tongue section under 20x magnification from a Phox2b-tdtomato reporter mouse to demonstrate proximity but no overlap in sensory and sympathetic fiber types.

Given evidence of anxiety and depression-like phenotypes in HNSCC patients, we sought to assess the potential contribution of these behaviors in tumor-bearing mice using the open field test (OFT) in tandem with the AnyMaze analysis software. OFT measures animal movement patterns in response to a novel, brightly lit environment, specifically preference for the center versus the periphery of the area. Anxiety and depressive-like behaviors are consistent with reduced exploration and increased time spent in the periphery (thigmotaxis), decreased locomotion, and increased immobility (*41*). Both MOC1 and MOC2-bearing mice at peak tumor burden exhibited significantly reduced travel distance and time spent in the center of the chamber compared to sham controls, indicative of increased anxiety-like behavior; changes in OFT movement measures were similar between MOC1 and MOC2 tumor mice **(Supp 1I-K)**. Pain can exacerbate this anxiety-driven avoidance; freezing behavior, defined as halted movement except for breathing for ≥3s, is a common defensive behavior in rodents when faced with aversive stimuli such as breakthrough spontaneous pain (*42, 43*). Notably, MOC2 tumor mice displayed significantly elevated freezing behavior (**Fig 2H,I**) that was reversible with analgesics (**Supp 1L**). Similar to MGS, there was no increase in freezing behavior observed in MOC1-bearing mice (**Fig 2I**) indicating spontaneous pain phenotype was specific to the MOC2 model. Lastly, there were no differences between the sexes in freezing behavior at PID12 (**Supp 1M**).

Collectively, these data suggest that the MOC2 tongue tumor model better recapitulates pain experienced by the HNSCC patient population reporting both functional and spontaneous/ongoing pain. To evaluate SNS hyperactivity that has also been denoted in HNSCC patients (*19*), we sought to quantify NE in MOC2 tumor-bearing and sham mice at peak nociceptive behavior (i.e. PID12). We were unable to separate enough plasma from tumor-bearing mice to isolate platelets for ELISA measurement, as done in the clinical population. Instead, we used mass spectrometry to detect and calculate the relative NE concentration in plasma and tumor tissue from tumor-bearing and sham mice. There was a significant increase in tongue tissue NE concentration from MOC2 mice compared to sham (p=0.047), but no change in mouse blood plasma (**Fig 2J**). We have also previously demonstrated a significant increase in CGRP protein in MOC2 tumor tissue compared to sham(*6*). These data suggest increased tumoral sympathetic and sensory signaling in the TME. Further examination of tongue tissue from both sham and MOC2 tumor-bearing mice identified potential gross anatomical changes in peripheral innervation over time. The anterior two-thirds of the tongue is densely innervated by sensory fibers primarily via the lingual nerve (i.e. a branch of the trigeminal mandibular nerve V3). Sympathetic postganglionic neurons (SPGNs) associated with the lingual nerve originate from the superior cervical ganglion (SCG) and reach their targets via a plexus on the facial artery. This dense innervation within a confined anatomical space likely contributes to the heightened pain reported by tongue cancer patients compared to those with other cancer types (*44*). To visualize peripheral innervation, we optically cleared 300µm tongue sections and stained them with the peptidergic sensory marker, CGRP, and the sympathetic marker, TH (**Fig 2K**). In sham mice, CGRP^+^ and TH^+^ immunofluorescence revealed dense nerve networks, with the prominent deep lingual artery running medially through the tongue with the lingual nerve alongside branching toward the epithelium (**Fig 2K**). By PID9, a large tumor mass was accompanied by CGRP^+^ nerve fiber encasement and clear disruption of the lingual nerve structure and deep lingual artery (**Fig 2K**). This suggests that tumor progression may be inducing nerve injury, alongside displacing and remodeling existing nerve structures (i.e. axonal sprouting). Using a *Phox2b*-Tdtomato reporter mice reporter mouse to identify SPGNs, magnification at 20x in 20µm sagittal tongue sections demonstrates sympathetic and sensory innervation in close proximity, evidenced by non-overlapping Tdtomato and CGRP-immunofluorescent fibers in the MOC2 TME (**Fig 2L**).

### 2.3 Oral Cancer Drives a Nerve Injury Response in Tongue-Innervating Sensory Neurons

Given the potential disruption of lingual nerve visualized in tumor sections particularly at later stages of tumor growth (PID9 and PID12), we sought to determine the extent of cancer-induced nerve injury (CINI) occurring in tongue innervating sensory and sympathetic neurons. Initial qualitative western blot analysis revealed an over two-fold increase in glial fibrillary acidic protein (GFAP) protein expression, a marker of Schwann cell activation/switch towards non-myelinating Schwann cells associated with nerve injury (*45, 46*), in MOC2 tongue homogenate compared to sham (**Fig 3A,B, Supp 2A**). During nerve injury Schwann cells have been demonstrated to dedifferentiate into a non-myelinating repair phenotype with migration towards the epineurium and surrounding tissue to aid in nerve regeneration response(*47–50*). To confirm the GFAP expression was related to nerve innervation near the tumor site, we stained 20µm sagittal tongue sections and found robust GFAP-immunofluorescence (IF) localized around and within the lingual nerve and lingual artery adjacent to the MOC2 tumor mass as well as at the tongue tip, compared to sham and MOC1 tongue sections (**Fig 3C,D, Supp 2B,C**). We confirmed GFAP-IF specificity in Schwann cells using Sox10 staining in sham and MOC2 tumor tissue serial sections and in a sciatic nerve ligation model (**Suppl Fig 3-4**); while Sox10-IF is evident in sham tissue, GFAP-IF is not present. These data support the conclusion that GFAP-IF is a marker of injury-induced activation. Peripheral nerve injury is associated with upregulation of transcription factors, *Atf3* and *Sox11* (*51, 52*). Whole ganglia real time qPCR from sham, MOC1 and MOC2 tumor bearing mice revealed significant increases in *Atf3* and *Sox11* relative expression at PID12 in TG from MOC2 mice compared to sham mice; there was no difference between MOC1 and sham mice (**Fig 3E, Supp 5A).** Unlike the peripheral sensory ganglia, there was no injury gene profile detectable in sympathetic ganglia (SCG) (**Fig 3E**) suggesting the GFAP staining was due to injury in sensory axons, not sympathetic. Given that the TG has vast heterogeneity in innervation targets and contains multiple cell types capable of expressing these target genes, including satellite glia and immune cells (*53, 54*), we assessed tongue-innervating neuron-specific expression using retrograde tracing and single cell qPCR. Consistent with whole ganglia data, there was a significant upregulation of relative *Atf3* gene expression at the single cell level in tracer positive neurons from MOC2 mice compared to sham, but no difference detected in tracer-positive neurons from MOC1 mice (**Fig 3F,G**). Immunostaining in TG and SCG sections confirmed a significant increase in ATF3 protein in tongue innervating tracer positive TG neurons from MOC2 mice compared to sham and MOC1 mice, but not in tracer positive SCG neurons (**Fig 3H,I**).

**Figure 3:**
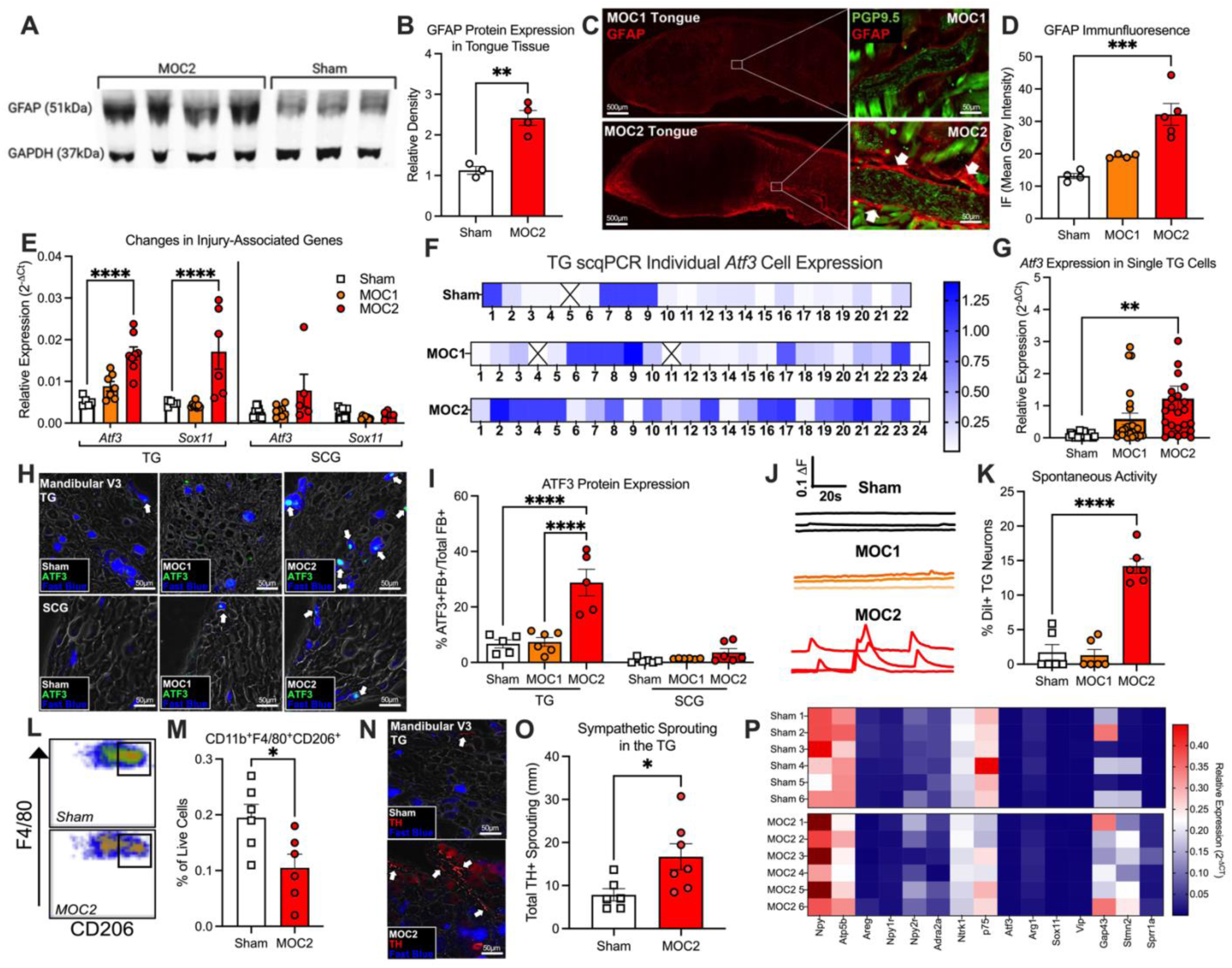
Oral cancer drives an injury response and plasticity in tongue-innervating sympathetic and sensory neurons: (A–B) Representative western blot and qualitative analysis of GFAP immunoreactivity in homogenized tongue tissue from MOC2 tumor-bearing and sham mice. GAPDH was utilized as the loading control. n=1-2/sex/group, Independent T-test, **p<0.01. (C–D) Immunofluorescence of GFAP in sagittal tongue section. Representative 4x (left) and 20x (Right) images of 20µm sections from MOC1 and MOC2-tumor bearing mice. Inset: 20x sagittal section of nerve bundle. 4x magnification highlights diffuse GFAP staining (red) and 20x magnification highlights increased immunofluorescence surrounding the lingual nerve in sections co-stained with pan-neuronal stain, PGP 9.5 (*127*), in MOC2 sections with minimal staining in MOC1 tongue tissue. White arrows indicate positive GFAP staining around nerve border. Quantification of relative immunofluorescence intensity (Mean Grey Intensity) through ImageJ was calculated relative to total section area in 4 tongue sections per mouse using 4x images from n=2-3/sex/group, One Way-ANOVA, ***p<0.005. (E) Gene expression analysis of injury-related genes in bilateral trigeminal ganglia (*93*) and superior cervical ganglia (SCG) from sham, MOC1 and MOC2 tumor-bearing mice. Relative gene expression of *Atf3* and *Sox11* were calculated following normalization to housekeeping gene *Gapdh*. n=3-4/sex/group, One-way ANOVA within tissue and gene comparisons, ****p<0.0001. (F–G) Single-cell qPCR of *Atf3* expression in retrogradely labeled tongue-innervating TG neurons. Gene expression was calculated as relative mRNA for each cell for each gene. *Gusb* was used as an internal control. Data are presented as heatmap for each cell. Boxes containing an X indicate no detectable expression (CT value > 38) for that cell (F). The relative expression distribution across treatment group was compared using Kruskal-Wallis test (G). n=22-24 neurons/group, **p<0.01 (H–I) ATF3 immunostaining in retrograde labeled (Fast Blue, FB) neurons from TG and SCG. Representative immunostaining of ATF3 protein in tracer positive (blue) neurons in 12µm TG sections (Top) and 10µm SCG sections (bottom) from sham (left), MOC1 (Middle), and MOC2 (Right) mice (H). Quantification of ATF3+FB+ neurons was calculated as a percentage relative to the total number of FB+ across 9 sections per mouse. n=2-3/sex/group, Independent T-Test within tissue type, ****p<0.0001. (J-K) Spontaneous calcium transients using Fura-2AM calcium imaging. Representative traces of non-evoked calcium transients in dissociated tracer-positive TG neurons from sham (top, black), MOC1 (middle, orange), or MOC2 (bottom, red) tumor-bearing mice. Spontaneous activity was defined as transients with a peak greater than 20% of baseline during a 60 second recording. n=3/sex/group, One Way-ANOVA, ****p<0.0001. (L–M) Flow cytometric analysis of dissociated trigeminal ganglia cells from sham and MOC2 mice at PID12. Representative scatter plots of CD11b^+^F4/80^+^CD206^+^ macrophages as a percent of live trigeminal cells. n=3/sex/group, Independent T-test, *p<0.05. (N-O) Tyrosine hydroxylase (TH) immunostaining in TG of Phox2b-Tdtomato reporter mice. Representative TH staining (red) surrounding tracer positive neurons (Fast Blue, blue) in V3 trigeminal branch in 12µm TG sections. Quantification of TH+ fiber density as measured by manual tracing of fiber tracks in ImageJ across ≥4 sections per mouse revealed elevated sympathetic fiber density in MOC2-bearing mice compared to sham. n=3-4/sex/group, Independent T-Test, *p<0.05. (P) Gene expression analysis of neurotransmission, injury and sprouting associated genes in bilateral whole SCG. Gene expression was calculated as relative expression for each sample for each gene. *Gapdh* was used as an internal control for normalization. Data are presented as heatmap for each gene and separated by treatment type (Sham, top; MOC2, bottom). n=3/sex/group.

ATF3 protein in tongue-innervating TG neurons suggests CINI and a neuropathic pain component in the MOC2 mouse model. Additional pathological hallmarks of neuropathic pain following peripheral sensory nerve injury are spontaneous neuronal activity (*55, 56*) as well as macrophage infiltration (*57*) and sympathetic postganglionic axonal sprouting (*58, 59*) into peripheral sensory ganglia. To functionally confirm CINI through activity, we used Ca^2+^ imaging to quantify spontaneous transients in acutely dissociated tracer positive tongue-innervating TG neurons in the absence of stimulation. While spontaneous activity was rare in tracer positive TG neurons from sham and MOC1 mice, approximately 15% from MOC2 mice displayed spontaneous Ca²⁺ transients at baseline (**Fig 3J,K**), consistent with nerve injury-induced hyperexcitability. We did not observe a size difference between the trigeminal neurons that were spontaneously active and those that were not. We also assessed macrophage activation and infiltration into TG using multi-color flow cytometry. Bilateral whole ganglia were processed from sham and MOC2 tumor-bearing mice at PID12. There was no significant difference in the frequency of CD11b^+^F4/80^+^ macrophages in the TG of tumor-bearing mice compared to sham (**Supp 5B,C**). However, the frequency of CD11b^+^F4/80^+^CD206^+^ M2 macrophages was significantly reduced in MOC2 tumor-bearing mice compared to sham (**Fig 3L,M**,), suggesting a phenotypic switch to proinflammatory M1 tissue macrophages in the TG of tumor bearing mice. Lastly, we performed TH immunostaining in TG tissue sections from retrograde labeled *Phox2b*-Tdtomato reporter mice to identify sympathetic innervation in the mandibular branch adjacent to tongue-innervating cell bodies. There was significant increase in total Tdtomato^+^TH^+^ fiber density in TG sections from MOC2 mice compared to sham (**Fig 3N,O**). Given that *Phox2b* lineage is also expressed in motor neurons, these data were further validated using a *Npy*-GFP reporter mouse; however, there was only a trend in increased innervation suggesting additional subpopulations of postganglionic sympathetic neurons may be involved (**Supp 5D,E**).

Evidence suggests that sympathetic postganglionic neurons can undergo plasticity in the context of disease (*60, 61*). Given that SCG neurons did not exhibit signs of injury but did demonstrate sprouting, we further investigated evidence of plasticity within the SCG. Transcriptomic profiles were assessed using whole ganglia PCR to evaluate genes associated with neurotransmission, activity, and target innervation. We found an upregulation of *Npy* as well as differential expression of NPY receptors, *Npy1r* and *Npy2r*, and autoreceptor gene *Adra2a* all thought to modulate NE and NPY neurotransmission (*62–65*) (**Fig 3P, Supp 5F**). While there were no changes in nerve injury genes *Atf3*, *Gal*, *Arg1*, or *Vip*, there was a significant upregulation of genes associated with regeneration and spouting, *Gap43*, *Stmn2* and *Sprr1a*. Lastly, nerve growth factor receptors, *Ntrk* and *p75*, known to regulate SPGN firing (*66*) were also differently regulated (**Fig 3P, Supp 5F**). To evaluate tumor-induced changes in SCG neuron activity, we evaluated acetylcholine (ACh, the predominant preganglionic input driver) evoked Ca^2+^ responses in dissociated SCG neurons from sham and MOC2 mice. There was a significant shift in the percentage of responsive SCG neurons to 3 and 30µM Ach as well as a significant increase in the magnitude of the Ach-evoked transient. Furthermore, KCl-evoked depolarization transients were also significantly larger in SCG neurons from MOC2 mice, suggesting changes in voltage gated Ca^2+^ channel expression (**Supp 5G-J)**. Together these data demonstrate MOC2 tumor-induced sympathetic sensitization, and a subsequent shift in sympathetic tone through the loss of *Adra2a* autoreceptor control.

### 2.4 MOC2 Oral Cancer-Induced Adrenergic Sensitivity via Alpha1 Adrenergic Receptors in Tongue Innervating Sensory Neurons

Previous studies in peripheral neuropathic pain models have posited that sensory nerves can acquire adrenergic sensitivity following peripheral nerve injury, defined as direct NE-evoked activity (*67–69*). Two mechanisms have been proposed for the development of adrenergic sensitivity: (1) upregulation of α1-adrenergic receptors (α1-ADRs), which are coupled to excitatory Gq-type GPCRs (*70–72*); or (2) a functional switch in α2-adrenergic receptors (α2-ADRs), coupled to inhibitory Gi proteins, to modulate N-type calcium channels post-injury (*73*). We have previously demonstrated that tongue-innervating TG neurons express genes for all adrenergic receptor subtypes with the exception of *Adrb3* (*74*). We first aimed to assess whether oral cancer progression evoked a shift in adrenergic receptor expression in tongue-innervating TG neurons. Single cell PCR in tongue innervating TG neurons from sham, MOC1 and MOC2 mice was used to probe for all ɑ- and β-adrenergic subtypes; RNA from 65 individual tracer-positive neurons (n=19-24 neurons across 3-4 mice per treatment group) revealed that all neurons contained detectable gene expression for at least 1 adrenergic receptor subtype. Tongue innervating neurons from sham animals exhibited the highest expression of ɑ2-adrenergic receptor subtypes, with ɑ1-subtypes showing the lowest expression levels (**Fig 4A,B**). While neurons isolated from MOC1 mice displayed a similar proportion and mean expression pattern to sham controls, there was a shift in the proportion and relative expression of α-ADRs in neurons from MOC2 mice. There was an increase in the number of tracer positive neurons expressing *Adra1d* as well as a small increase in mean expression. The number of tracer positive *Adra2b* and *Adra2c* expressing neurons remained similar, but mean expression of each gene was significantly decreased in neurons from MOC2 mice (**Fig 4A,B**).

**Figure 4:**
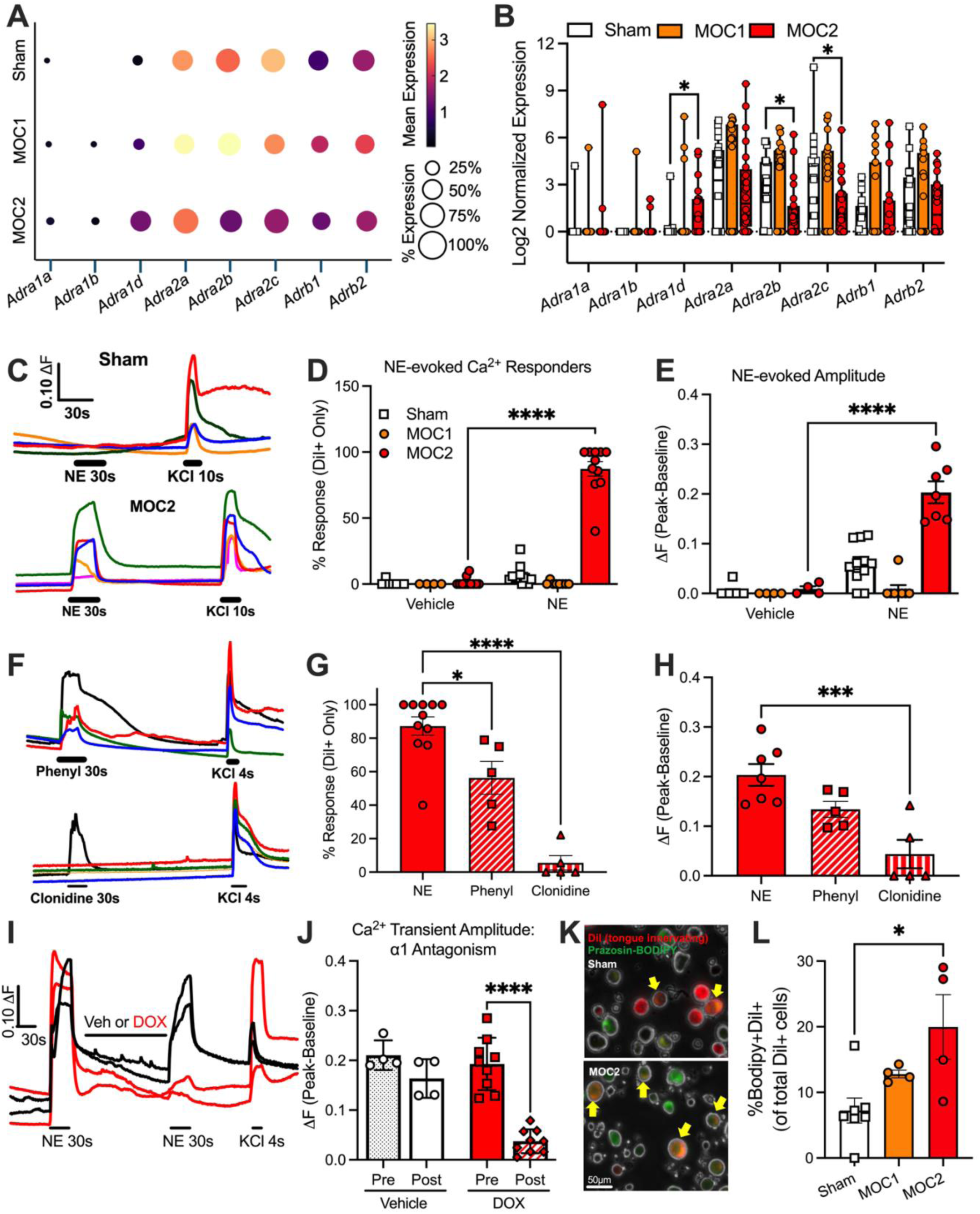
Oral cancer-induced adrenergic activity via α1 adrenergic receptors in tongue-innervating sensory neurons. (A–B) Single-cell qPCR analysis of tongue-innervating (tracer-positive) trigeminal ganglion neurons from sham, MOC1, and MOC2 mice revealed altered adrenergic receptor gene expression profiles. n=19-24 cells/group, Two-way ANOVA, Cancer by gene interaction, *p<0.05. (C–E) Calcium imaging of dissociated tongue-innervating (DiI+) neurons using Fura-2AM. Representative calcium transients evoked by 10 µM norepinephrine (NE) and depolarization evoked by 30mM KCl (C). Quantification of the percentage of DiI+ neurons responsive to both NE and KCl (D) as well as the amplitude (peak – baseline) of the NE-evoked responses (E). n=5-6/sex/group, Two-Way ANOVA, Cancer by treatment interaction, ****p<0.0001. (F-H) Adrenergic agonist pharmacology in dissociated tongue-innervating (DiI+) neurons using Fura-2AM. Representative calcium transients evoked by 10µM phenylephrine (α1 agonist), 10µM clonidine (α2 agonist) and 30mM KCl application (F). Quantification of the percentage of DiI+ neurons responsive to 10µM NE, phenylephrine, and clonidine (G) as well as the amplitude of the evoked response (H). n=2-6/sex/group, One-Way ANOVA, *p<0.05, ***p<0.005, ****p<0.0001 (I-J) Adrenergic alpha 1 (α1) antagonist pharmacology in dissociated tongue-innervating (DiI+) neurons using Fura-2AM. Representative calcium transients evoked by 10 µM norepinephrine (NE) before and after 2-minute application of vehicle (0.01% DMSO in normal bath, black) or 10µM doxazosin mesylate (DOX, red), a selective α1-adrenergic receptor antagonist (I). Quantification of NE-evoked transient amplitude prior to (pre) and after (post) DOX treatment. n=2-4/sex/group, Two-way ANOVA, Treatment by time interaction, ****p<0.0001. (K-L) Protein expression of α1-adrenergic receptors in TGNs was confirmed using prazosin-BODIPY binding. n=2-3/sex/group, One-Way ANOVA, *p<0.05.

**Figure 5:**
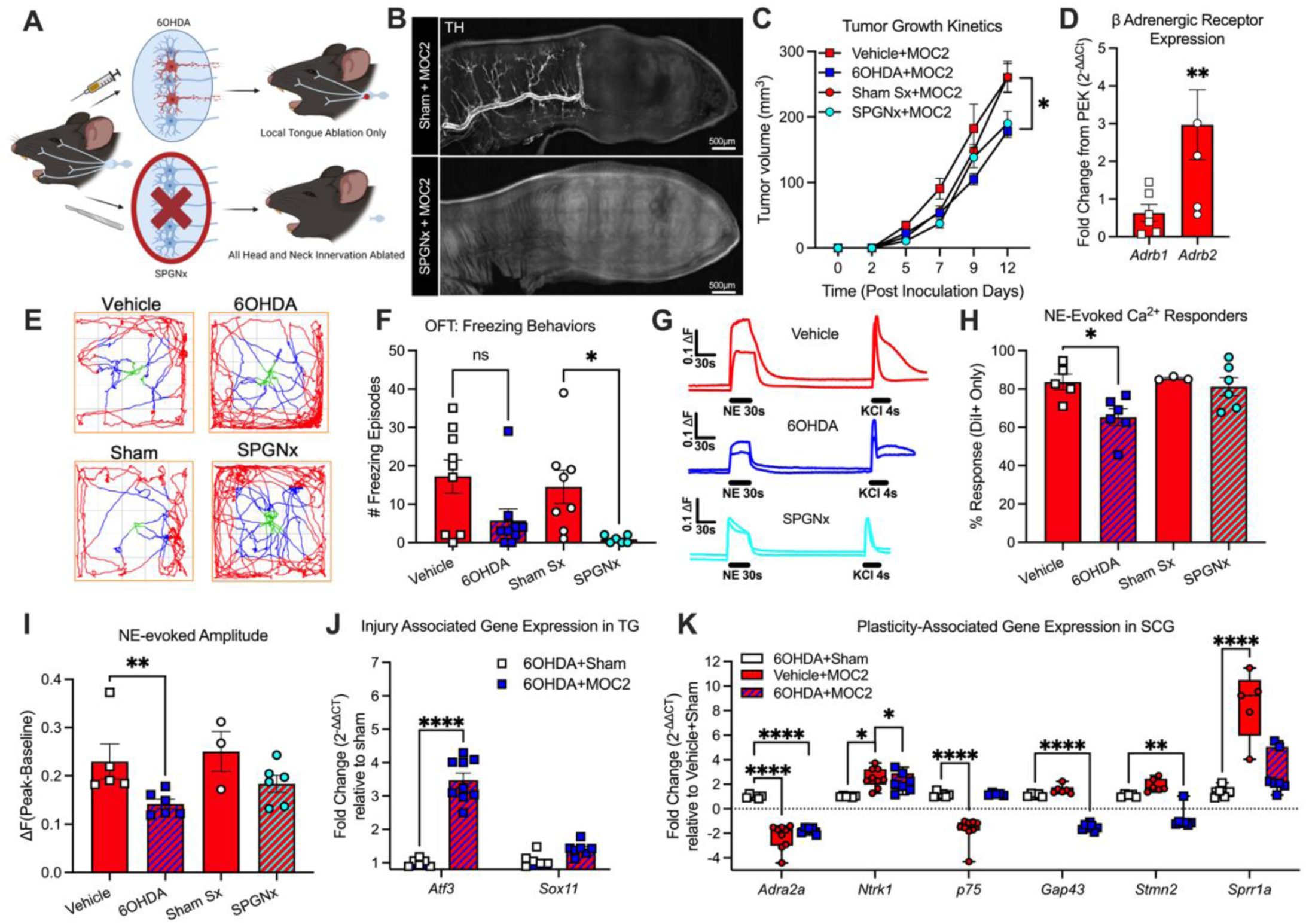
Loss of local sympathetic innervation attenuates tumor-induced pain behavior but not adrenergic sensitivity. (A) Schematic of two sympathectomy methods. Top: local chemical sympathectomy via anterior and posterior tongue injections of 6-hydroxydopamine (6OHDA) to ablate local sympathetic fibers. Bottom: surgical superior cervical ganglionectomy (SPGNx) eliminates sympathetic innervation to the head and neck region. (B) Representative tyrosine hydroxylase (TH⁺) immunofluorescence staining of 300µm sagittal tongue sections from sham and SPGNx mice, confirming successful loss of sympathetic innervation 1 week following surgery. (C) Tumor growth kinetics of MOC2-bearing that underwent 6OHDA or SPGNx treatments compared to sham and vehicle tumor bearing controls by PID12. n=3-5/sex/group, Two-Way ANOVA, Treatment by time interaction, *p<0.05. (D) Beta1-Adrenergic (*Adrb1*) and Beta2-Adrenergic (*Adrb2*) receptor gene expression in MOC2 cell line compared to non-tumorigenic primary epithelial keratinocytes (PEK). n=4-6 passages/group, Independent T-test of delta-CT values compared to PEK, *p<0.05. (E-F) Open Field Test (OFT) analysis of freezing episodes. Representative track plots and quantification of episodes across sympathectomize and sham/vehicle MOC2 mice. n=3-5/sex/group, Independent T test within methodology (i.e. 6OHDA versus vehicle; SPGNx versus sham), *p<0.05. (G-I) Calcium imaging of tongue-innervating TGNs to test for potential changes in acquired adrenergic sensitivity following sympathectomy. Representative calcium transients in response to 10µM norepinephrine (NE) and depolarization evoked by 30mM KCl (G). Quantification of the percentage of tracer-positive neurons responsive to both NE and KCl (H) as well as the amplitude (peak – baseline) of the NE-evoked responses (I). n=3-6/group, One-Way ANOVA, *p<0.05, **p<0.01. (J) Gene expression analysis of injury-associated genes *Atf3* and *Sox11* in TG from sham and MOC2 tumor-bearing mice following chemical sympathectomy. n=3-4/sex/group, Independent T-Test, ****p<0.0001. (K) Gene expression analysis of plasticity-associated genes in bilateral SCG from mice that received chemical (6OHDA) sympathectomy prior to MOC2 or sham inoculation. Changes in expression were determined by fold change from SCG from mice that received vehicle+sham treatment. Neurotransmission associated gene (*Adra2a, Ntrk1, p75*) and axonogenesis associated genes (*Gap43, Stmn2, Sprr1a*). Two-way ANOVA, Cancer by treatment interaction within gene, *p<0.05, **p<0.01, ****p<0.0001.

Due to primarily inhibitory α2 receptor expression, adrenergic activation on sensory neurons, including tongue-innervating trigeminal neurons^27^, does not result in depolarization or neuronal firing (*75–77*). However, given that the expression data suggested a phenotypic switch to an excitatory Gq/11-coupled α1 adrenergic receptor profile, we utilized Ca^2+^ imaging with Fura-2AM as Ca^2+^ indicator to investigate a potential acquisition of calcium flux in response to NE (10µM) or vehicle (0.001% HCl) across groups. There was a significant interaction between pharmacology and tumor type regarding the percentage of NE-responsive tongue afferents as well as the amplitude of evoked response. Consistent with previous results, we found that neither vehicle nor NE evoked a Ca^2+^ transient greater than 20% of baseline Ca^2+^ in most tracer-positive neurons from sham mice; only 6.8% of tongue afferents responded with an average peak only 6% greater than baseline (**Fig 4C-E**). However, in neurons from MOC2 mice, NE evoked a substantial Ca^2+^ transient; 91.2% of tongue afferents responded (p<0.0001, **Fig 4C,D**) with an average peak 115% greater than baseline (p<0.0001, **Fig 4E**). Of note, approximately 20% of non-tracer positive cells also exhibited a response to NE (**Supp 6A**). To test receptor-specific contributions, we evaluated Ca²⁺-evoked transients using selective α1 and α2 agonists, phenylephrine and clonidine, respectively, as well as selective β1 and β2 agonists, xamoterol and salbutamol, respectively. Less than 1% of tongue afferents responded to 10µM clonidine, whereas over 60% responded to 10µM phenylephrine (F(2,18)=36.76, p<0.0001) with peak amplitudes significantly elevated above baseline levels (p<0.0008, **Fig 4F-H**). Additionally, doxazosin mesylate, a selective α1 antagonist, significantly reduced NE-evoked Ca²⁺ transients compared to vehicle controls (**Fig 4I,J**). Neither β1 or β2 agonist elicited Ca^2+^ transients in TG from MOC2 tumor-bearing mice (**Supp 6B,C**). Alpha 1 receptor protein expression was subsequently confirmed in TG neuron cultures using highly selective α1-adrenergic receptor antagonist Prazosin conjugated to a boron-dipyrromethene (BODIPY) fluorophore (p<0.0195, **Fig 4K-L**). Taken together, these findings suggest that MOC2 tumor progression promotes adrenergic receptor plasticity in tongue-innervating TG neurons, particularly through upregulation of α1-ADRs, potentially contributing to heightened sensory neuron activity and transduction of nociceptive signaling towards the central nervous system.

**Figure 6:**
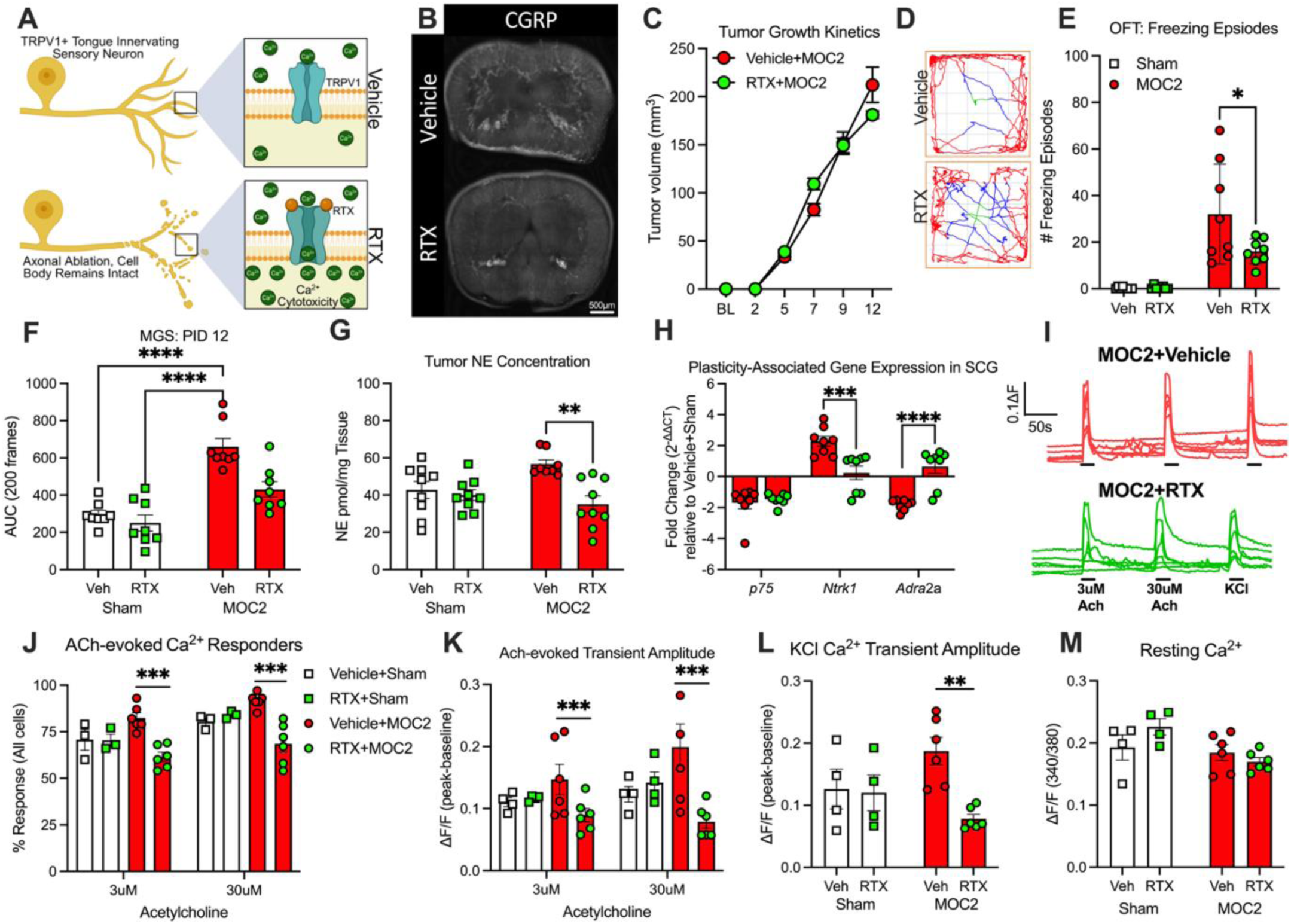
Loss of local TRPV1^+^ sensory innervation reduces tumor-induced sympathetic dysregulation. (A) Schematic of RTX mediated denervation. Local RTX administration into the tongue drives cytotoxicity through TRPV1 cation channel expressed on tongue-innervating trigeminal axons resulting in local denervation. TRPV1^+^ cell bodies housed in the trigeminal ganglia remain intact. (B) Representative immunostaining of CGRP-positive nerve density in 300µm thick coronal sections from vehicle and RTX treated tongues from MOC2 mice at PID12. Sections used were directly posterior to tumor-containing tissue and confirms a reduction in sensory innervation 3 weeks after RTX treatment. (C) Tumor growth kinetics in vehicle and RTX-treated MOC2 tumor bearing mice measured by caliper every 3 days, with no change in tumor growth by PID12. n=4/sex/group Two-way ANOVA, Treatment by time interaction, p>0.05 (D-E) Spontaneous freezing behavior was quantified using the Open Field Test (OFT). Representative track plots (D) and quantification of total number of freezing episodes per mouse across groups (E). n=3-4/sex/group, Two-Way ANOVA, Cancer by RTX treatment interaction, *p<0.05. (F) Mouse grimace score (MGS) area under the curve (AUC) analysis for grimace behavior across groups. n=4/sex/group, Two-way ANOVA, Cancer by treatment interaction, ****p<0.0001 (G) Norepinephrine (NE) concentration in tongue tissue from vehicle and RTX treatment sham and MOC2 tumor bearing mice at PID12 measured by mass spectrometry. n=3-5/sex/group, Two-Way ANOVA, Cancer by treatment interaction, **p<0.01. (H) Gene expression analysis in bilateral SCG from MOC2 tumor bearing mice pretreated with either vehicle or RTX. n=4/sex/group, Two-Way ANOVA, Treatment by Gene interaction, ***p<0.005, ****p<0.0001. (*128*) Percent of acetylcholine (3uM, 30uM)-responsive neurons and amplitude of evoked response in dissociated SCG from vehicle and RTX-treated sham and MOC2 mice; L) Amplitude of depolarization-evoked transient. M) resting calcium levels. n=2-3/sex/group, Two-Way ANOVA, Cancer by treatment interaction, **p<0.01, ***p<0.005.

### 2.5 Sympathetic Denervation Reveals Adrenergic Contribution to Tumor Growth and Tumor-induced Nociceptive Behavior

Our data suggests OSCC-associated dysregulation of sympathetic efferent activity coupled with α1-mediated adrenergic sensitivity in sensory neurons. We sought to determine whether this mechanism contributes to tumor-associated nociceptive signaling *in vivo* through targeted sympathetic denervation. Localized 6-hydroxydopamine (6OHDA) injections to chemically ablate tongue sympathetic postganglionic nerve fibers or surgical SCG sympathectomy (SPGNx) two weeks prior to tumor inoculation resulted in a marked reduction in TH^+^ fibers in the tongue compared to sham-operated or vehicle-injected controls, while leaving other nerves intact (**Fig 5A,B, Supp 7A,B**). It is well-established that the SNS can influence tumor progression (*1, 78*); direct modulation of tumor proliferation by sympathetic catecholamines has been demonstrated in many solid tumor lines (*79*), including HNSCC (*1, 80*). Thus, not surprisingly, we found that both denervation strategies suppressed tumor growth and reduced weight loss in MOC2 mice by PID12 (**Fig 5C**, **Supp 7C**). Assessment of adrenergic receptor expression in mouse oral cancer cell lines confirmed that both lines expressed genes for β-ADRs; however, there was 3-fold more *Adrb2* expression in the non-immunogenic MOC2 cell line compared to primary keratinocyte cell line (PEK) (**Fig 5D**) suggesting sympathetic postganglionic input could impact tumor proliferation directly via beta adrenergic receptor expression on the tumor cells thus loss of sympathetic innervation may slow growth. Additionally, evaluation of alpha-adrenergic receptor expression previous underappreciated in HNSCC cell lines revealed disparate expression of α1 and α2 subtypes; indolent immunogenic MOC1 cells contained higher relative expression of inhibitory α2-ADR subtypes compared to aggressive non-immunogenic MOC2 **(Supp 7D-F)**.

**Figure 7:**
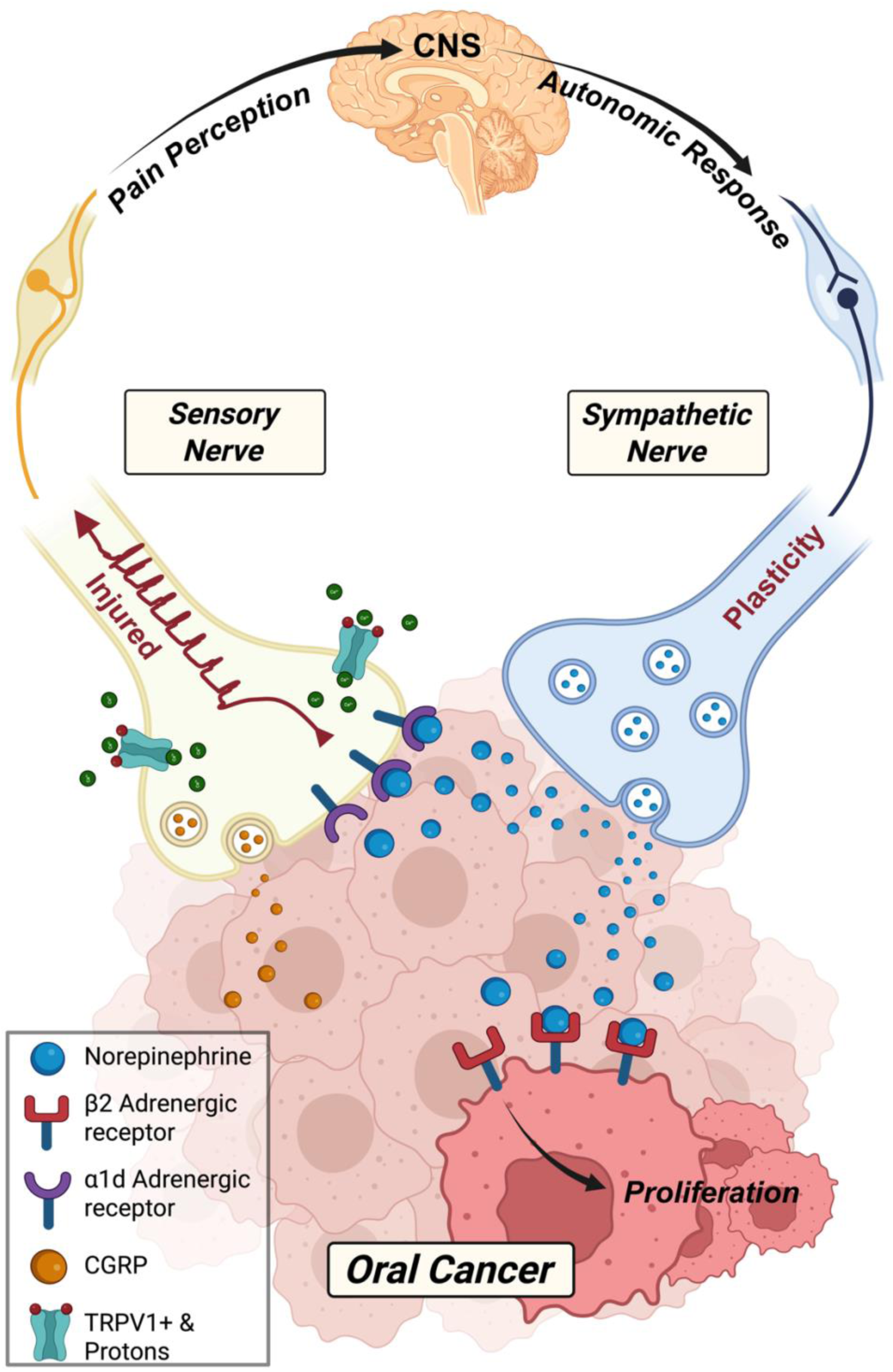
Sympathetic Plasticity and Sensory Injury Create a Maladaptive Feed-Forward Loop Driving Oral Cancer Nociception: Schematic representation of cancer-induced nerve injury (CINI) in oral squamous cell carcinoma. Tumor growth injures lingual sensory nerves, leading to increased release of calcitonin gene-related peptide (CGRP, orange). In parallel, sympathetic postganglionic neurons undergo plasticity, sprouting into the tumor microenvironment and releasing norepinephrine (blue). Sensory neurons acquire adrenergic sensitivity via upregulation of α1d-adrenergic receptors (purple), while tumor cells express β2-adrenergic receptors (red). This maladaptive crosstalk promotes spontaneous pain and enhances tumor proliferation, establishing a maladaptive feed-forward loop that integrates peripheral nerves with the central nervous system (CNS).

Next, we sought to measure nociceptive behavior directly in the sympathectomized tumor-bearing mice compared to intact. We evaluated freezing and grimacing behaviors using the OFT and MGS, respectively, at PID12. SPGNx-treated mice demonstrated a significant reduction in freezing behavior; however, MGS assessment was not possible due to surgical sympathectomy-induced ptosis, resulting in an eye artifact (*81, 82*). In 6OHDA-treated MOC2 mice, there was a trending reduction in both freezing behavior and facial grimacing compared to controls (**Fig 5E,F**, **Supp 7G-I**). These results suggest that loss of sympathetic postganglionic input results in decreased spontaneous pain-like behavior in mice. While grimacing behavior only exhibited a nonsignificant downward trend, this could be due to other tumor-secreted factors (i.e. TNFα(*1*)) that can also drive nociception as the sensory nervous system is still intact. However, open field freezing behavior indicated a robust impact of SPGN on spontaneous pain-like behavior.

We posited that slowed tumor growth in sympathectomized mice impacted neuronal plasticity and development of adrenergic sensitivity in sensory neurons. Thus, we performed Ca^2+^ imaging on tongue innervating TG neurons from sympathectomized MOC2 tumor bearing mice at PID12 (**Fig 5G**). There was a small decrease in percentage responsive to NE application as well as the amplitude of response in tracer positive TG neurons from 6OHDA treated mice, but no significant difference in NE-evoked Ca^2+^ transient in TG neurons from SPGNx treated mice compared to MOC2 mice with intact sympathetic innervation (**Fig 5H-I**). Further, whole ganglia qPCR of TG from 6OHDA-treated MOC2 mice revealed that the injury marker ATF3 was still upregulated compared to sham (**Fig 5J**), despite the reduced tumor growth in sympathectomized mice. Given that the sympathectomy results in loss of sympathetic tongue innervation prior to inoculation, we used real time qPCR to evaluate the SCG neuronal plasticity in 6OHDA-treated MOC2 mice compared to sham and intact MOC2 mice. Genomic analysis revealed that chemical sympathectomy prevented the tumor-induced increase in sympathetic sprouting genes *Gap43*, *Stmn3*, and *Sprr1a* as well as the loss in *p75*. However, surprisingly, 6OHDA did not prevent the tumor-induced downregulation of autoreceptor *Adra2a* expression or the increase in *Ntrk1* (**Fig 5K**). Collectively, these findings indicate that direct sympathetic signaling contributes to tumor growth and may influence tumor-evoked nociception through adrenergic-mediated sensory neuron plasticity. However, tumor-associated sympathetic input is not strictly required for sensory adrenergic plasticity or some changes in tumor-induced sympathetic dysregulation.

### 2.6 Loss of TRPV1+ Fibers Reduces Sympathetic Tone

Emerging evidence suggests a connection between sensory circuits in oral cancer and the modulation of central nervous system (CNS) nuclei associated with autonomic regulation (*9*). We hypothesize that in the MOC2 oral cancer model, sensory input from the TME drives increased sympathetic tone, creating a feedback loop wherein elevated NE levels exacerbate spontaneous pain. To test this hypothesis, we selectively ablated a peptidergic subpopulation of tongue-innervating sensory neurons prior to tumor inoculation using resiniferatoxin (RTX), a toxin specific to the TRPV1 channel that when administered in the tissue results in reversible sensory denervation (*83*) (**Fig 6A, Supp Fig 8A**). We have previously demonstrated a role for TRPV1-expressing sensory neurons in oral cancer-associated pain (*84*). CGRP neuropeptide has high co-expression in TRPV1⁺ neurons (*85, 86*) and an established role in modulating tumor growth via the CGRP-RAMP1 signaling pathway (*6, 7, 87, 88*). Immunostaining of optically cleared 300 µm coronal tongue sections confirmed successful RTX-mediated sensory neuron ablation, demonstrated by substantial reduction in CGRP⁺ fiber density in RTX-treated tissues compared to vehicle (0.9% NaCL, 0.25% Tween-80, 2mM ascorbic acid) controls (**Fig 6B**).

Consistent with findings from other non-immunogenic cell line studies(*7*), we found that RTX-mediated denervation prior to inoculation did not significantly impact tumor growth over the observed period in non-immunogenic MOC2 tumor-bearing mice (**Fig 6C**). However, spontaneous freezing episodes during the OFT were significantly reduced in RTX-treated MOC2 mice compared to vehicle-treated MOC2 mice accompanied by a significant decrease in grimacing behavior at PID12 (**Fig 6D-F**), suggesting a reduction in cancer-induced sensory input and spontaneous/ongoing pain. RTX- and Vehicle-treated sham mice did not exhibit any pain-like behaviors and MOC2-induced anxiety-like OFT behaviors remained unaffected by RTX treatment (**Supp Fig 8B-G**).

We posited that sensory adrenergic sensitivity is driven by tumor-induced nerve injury. Thus, we investigated whether RTX-mediated sensory denervation, shown to drive ATF3 expression (*89–91*), induced adrenergic sensitivity in sham mice or attenuated its development in MOC2-tumor bearing mice. Using Ca^2+^ imaging in dissociated TG neurons across groups revealed that RTX ablation did not induce adrenergic sensitivity nor did it reduce the percentage of NE-evoked responses in tracer-positive trigeminal ganglion neurons (**Supp Fig 8H-J**). Therefore, together these data suggest that ablating a subset of sensory neurons partially mitigated nociceptive behaviors by reducing sensory input to the CNS but did not affect the development of adrenergic sensitivity.

To further test our hypothesis that sensory input from the TME is involved in sympathetic dysregulation, we used mass spectrometry to quantify tumoral NE concentration in RTX-denervated and intact MOC2 mice. Interestingly, we found a significant interaction between RTX denervation and cancer; RTX-denervation prior to inoculation blocked the elevated NE concentration in tumors from MOC2-bearing mice (**Fig 6G**), suggesting reduced sensory neuron input to the CNS may subsequently lower sympathetic output. Given that sympathetic denervation was unable to reverse components of the observed tumor-induced sympathetic neuron plasticity, we used qPCR to evaluate changes in SCG from vehicle- or RTX-treated MOC2 mice. While there were no differences in *p75* expression, RTX treatment blocked the downregulation of autoreceptor *Adra2a* and the upregulation of *Ntrk1* (**Fig 6H**). Functional testing using Ca^2+^ imaging in dissociated SCG neurons confirmed the loss of disrupted sympathetic tone following RTX treatment (**Fig 6I**). RTX significantly attenuated the previously observed increase in the percentage of neurons responsive to acetylcholine (Ach) at concentrations of 3 and 30 µM, as well as the tumor-induced increase in Ca^2+^ transient amplitude evoked by both Ach and KCl (**Fig 6J-L**). Resting calcium levels remained consistent across all groups (**Fig 6M**). These data suggest that diminished sensory neuron activity influences tumor-mediated sympathetic postganglionic neuronal plasticity and sensitization.

## 3.0 Discussion

Our findings provide compelling evidence that CINI in head and neck cancer promotes maladaptive crosstalk between the sympathetic and sensory nervous systems within the TME, contributing to the development of spontaneous, neuropathic-like pain and subsequent sympathetic dysregulation. We demonstrate that SPGNs undergo bidirectional plasticity in response to both tumor-derived signals and afferent sensory activity, highlighting a complex and dynamic interplay between peripheral neural circuits during oral cancer progression (**Fig 7**).

### Sympathetic Activity Correlates with Spontaneous Pain in OSCC Patients

Heightened SNS activity has been previously reported in HNSCC patients; Bastos et al found that sympathetic catecholamines NE and epinephrine are elevated in oral and oropharyngeal SCC patients compared to non-cancer patients (*19*). Building upon this foundation, our prospective observational study reveals a strong correlation between elevated circulating NE levels and patient-reported pain in OSCC. Stratification of pain into function-evoked and spontaneous revealed that spontaneous pain is an independent predictor of PNI, aligning with the neuropathic characteristics frequently observed in OSCC-related pain (*16*). While the hypothalamic-pituitary-adrenal (HPA) axis contributes to systemic NE levels, approximately 10–20% of circulating NE originates from the adrenal medulla (*92*), with the remainder released from sympathetic nerve terminals. Evidence from cancer patients further suggests that localized NE release at the tumor site may play a more dominant role in modulating tumor progression and pain than systemic HPA contributions (*93*), supporting the use of circulating NE as a proxy for SPGN neurotransmission, rather than generalized stress. While not possible with the current dataset, further characterization of the neuropathic pain components (*94*) with a more diverse patient population is warranted through specific questionnaires that improved phenotyping and stratification of pain in patients will help address the underlying etiology and enable more personalized therapeutic approaches. Additionally, retrospective and prospective evaluation of concomitant adrenergic pharmacology use with standard of care and its impact on pain is necessary for clinical translation of our findings.

### MOC2 Model Recapitulates Clinical Phenotypes

MOC2 tongue tumor growth recapitulated the patient-reported function-evoked and spontaneous/ongoing pain with oral SCC. However, elevated systemic (plasma) NE was not recapitulated and remained unchanged in tumor mice compared to sham control suggesting tumor progression does not induce a generalized stress response measurable by catecholamines. We posit that this lack in translation is likely due to methodology; circulating NE levels were measured in platelets isolated from 1mL of human plasma quantified by ELISA whereas mass spectrometry was used to quantify NE in 50uL of mouse plasma. While mass spectrometry is a more sensitive technique, fluctuations in plasma catecholamines are well documented (*29, 30*) and we were unable to isolate enough platelets from mouse plasma in tumor-bearing animals to use ELISA due to disease mediated weight loss. Given the data from cancer patients supporting a sympathetic-peripheral rather than stress-systemic interpretation of circulating NE, failure to detect a change in circulating NE in the mouse is likely a methodological issue. However, previous literature has also demonstrated that nude mice with tongue SCC failed to display anxiety-like behaviors as measured by elevated maze and lacked evidence for higher activity in anxiety-associated brain regions (*95*). Moreover, studies by Repasky and colleagues suggest that mice held in standard housing conditions are already experiencing a thermal stress and alleviating stress by shifting them from the hypothermic to thermoneutral conditions (*96*). Elevated NE was measured in the tumor tissue from mice compared to sham tongue tissue held at the same thermal conditions support the hypothesis that localized sympathetic activity contributes to tumor-associated nociceptive signaling.

### Cancer-Induced Nerve Injury Drives Sensory Neuron Plasticity

The anatomical convergence of lingual nerve and vascular structures in a restricted tongue compartment likely exacerbates vulnerability to tumor-induced injury and may underlie the intensity of spontaneous pain reported by patients with tongue cancer (*44*). Evaluation of injury through gross anatomical assessment, gene and protein expression, and functional activity all indicated that MOC2 tumor growth in the tongue established an injury phenotype in sensory but not sympathetic neurons. Injury in sensory neurons in the dorsal root system have been widely studied through sciatic nerve ligation and transection models establishing *Atf3* and *Sox11* as key transcription factors associated with sensory neuron injury (*51, 52*) along with macrophage expansion (*57*) and increased sympathetic innervation (*58, 59*) into the sensory ganglia. While sympathetic injury has been severely understudied, early evidence suggests *Atf3* is upregulated with sympathetic chain transection (*97, 98*). We did not detect significant changes in 4 different injury related genes at the SCG whole ganglia level; however, additional single cell evaluation is needed to fully appreciate the transcriptomic shifts in SPGN neurons during tumor progression. Nonetheless, tumor-induced sympathetic plasticity was evident, as MOC2-bearing mice showed increased TH⁺ fiber density in TG sections, suggesting sympathetic postganglionic axonal sprouting into the sensory ganglia; a hallmark of nerve injury pain states (*58*). Furthermore, several genes linked to sympathetic neurotransmission (*Nktr1*, *p75*, *Adra2a*), as well as sprouting genes (*Gap43*, *Sprr1a*) all demonstrated changes in the MOC2 model suggestive of sympathetic plasticity. Neurotrophic receptors, TrkA and p75, are proposed to have opposing effects on the firing properties of sympathetic neurons; activation of TrkA promotes tonic firing, while p75 promotes phasic firing (*66*). This suggests that the relative level of signaling through these receptors can regulate the neuron’s electrical activity and ultimately the functional output of the sympathetic nervous system (*66*).

### Acquisition of Adrenergic Sensitivity via α1-Adrenergic Receptor Plasticity

Sympathetic-to-sensory signaling at homeostasis likely serves a suppressive role in nociception, consistent with the well-documented analgesia observed during states of heightened autonomic activity (e.g., the “fight-or-flight” response) (*99*). Under homeostatic conditions, sensory neurons predominantly express the inhibitory α2-ADR receptor (*100–102*) and activation provides robust inhibition of TRPV1-mediated activity in healthy dorsal root ganglion neurons (*77, 102*). This study unveils a significant shift in adrenergic receptor subtype expression in the presence of tumor-induced nerve injury. Aggressive oral cancer progression promotes functional adrenergic sensitivity in tongue-innervating sensory neurons through upregulation of α1-ADRs. This acquired phenotype aligns with mechanisms previously described in neuropathic pain conditions, including peripheral nerve injury (*68, 69*) and complex regional pain syndrome, where enhanced sympathetic-sensory coupling contributes to persistent pain (*103*). Activation of α1-ADR receptors has been shown to mobilize intracellular calcium as well as activate calcium influx via voltage-dependent calcium channels (*104, 105*). Multiple mechanisms may contribute to the upregulation of α1-ADRs following nerve injury. Injury-induced stress signals, such as elevated intracellular calcium, reactive oxygen species, and proinflammatory cytokines, activate several intracellular signaling pathways, including CREB1, and MAPK pathways (*106–108*). CREB1 has been implicated in the transcriptional regulation of adrenergic receptor genes (*109*) and has been noted as being phosphorylated by alpha1 adrenergic receptor activation, suggesting a reciprocal relationship between CREB1 singling and adrenergic receptor activity (*110*). The MAPK pathway activation can lead to JNK mediated phosphorylation of c-Jun and increased expression of ATF3 (*111*). Together, along with other proteins in the Fos family, they can form the Activator Protein-1 (AP-1) transcription factor complex (*112*), which has been implicated in regulating adrenergic receptor expression(*113*). Additionally, the MAPK pathway can activate Sp1, a transcription factor specifically linked to the expression of α1-ADRs (*114, 115*).

### Sympathetic Signaling Promotes Tumor Progression and Modulates Spontaneous Pain Behavior

Beta adrenergic receptor expression is linked to tumor proliferation in many solid tumor types through direct tumor signaling and indirectly via the immune system (*75*). Targeted sympathetic denervation reduced MOC2 tumor growth, likely mediated by NE signaling directly on β-ADRs on cancer cells (*1, 79, 80*) given its non-immunogenic phenotype. We also acknowledge that MOC2 cells did express α1-ADRs which are characterized as excitatory GPCRs and thus it is possible that α1 activation on MOC2 through sympathetic input supports proliferation. Future investigation of α1 antagonism on oral cancer proliferation and migration is warranted. Importantly, the loss of sympathetic tone was also associated with decreased nociceptive behavior despite injury-associated *Atf3* remaining elevated in TG. Loss of sympathetic tone in oral cancer did not ablate all tumor-associated nociceptive behavior suggesting that while sympathetic signaling may contribute to the magnitude nociception, it is not strictly necessary for its development. This aligns with the possibility that CINI or even additional tumor-derived mediators, such as pro-inflammatory cytokines (e.g., TNF-α), may contribute and drive α1-ADR upregulation or facilitate a pro-nociceptive transcriptional program in sensory neurons. These findings support previous literature implicating SNS activity in promoting tumorigenesis and nociception in head and neck cancers (*1, 75, 76*).

### TRPV1⁺ Neuron Ablation Attenuates Sympathetic Plasticity and Spontaneous Pain

Previous cancer neuroscience studies have begun to test the hypothesis that nociceptive input is required for amplified sympathetic output via central integration in autonomic control centers (*7*). Our findings support this hypothesis with two key results. First, while targeted local ablation of TRPV1⁺ sensory neurons did not alter MOC2 tumor growth, a finding consistent with prior studies using similar non-immunogenic tumor models (*7*), there was a reduction in tumoral NE concentration as well as spontaneous pain-like behaviors, implicating a role for sensory neurons in driving elevated sympathetic tone. These results are consistent with establishing a feed-forward loop that exacerbates pain through elevated NE signaling and acquired adrenergic sensitivity (**Fig 7**). Second, molecular and functional analyses of the SCG revealed bidirectional modulation of sympathetic output. Loss of SPGN innervation into the TME through chemical sympathectomy was only able to rescue some of the tumor induced gene changes, most notably genes associated with peripheral sprouting. Interestingly, loss of TRPV1^+^ sensory innervation into the TME recovered SCG expression of the *Adra2a* autoreceptor and *Ntrk1* neurotrophic receptor. Further, SCG neurons from RTX-treated mice displayed significantly reduced responsiveness to preganglionic input (i.e. ACh) and depolarization, confirming that diminished sensory input reduces postganglionic sympathetic output. Given that the sympathetic signaling relies on a 2-neuron efferent system, we posit that afferent input is driving sensitization in preganglionic input resulting in plastic signaling within the sympathetic ganglia.

### Limitations of the Study

This study has several limitations that warrant consideration. First, while our analyses reveal significant correlations between psychological symptom burden and patient-reported pain, the sample size to detect pain and NE as predictors of PNI were underpowered. However, these data highlight patient-reported pain and sympathetic signaling as potential biomarkers and suggest that further exploration is warranted. Second, although we demonstrate reciprocal plasticity between sympathetic and sensory neurons, our study does not comprehensively address the role of the tumor vasculature or neuroimmune crosstalk in mediating these interactions. The tumor microenvironment is highly vascularized, and emerging evidence suggests that blood vessels serve as critical conduits for nerve-endothelial communication, with implications for both tumor progression and pain signaling. Our optical clearing and immunostaining approaches captured nerve-vessel proximity, particularly along the lingual artery, but did not assess how vascular remodeling during tumor growth might influence nerve injury or sympathetic sprouting. Additionally, denervation strategies (*7, 116*) and cancer-induced nerve injury (*22, 117*) can impact the tumor-associated immune response thus investigation of the downstream immunological consequences of this sympathetic-sensory neural circuit is warranted. Lastly, while our findings suggest that α1-adrenergic receptor antagonists may have therapeutic potential, translation to clinical practice will require additional preclinical validation, including testing in genetically engineered models or carcinogen-induced models are needed to better elucidate the dynamic and longitudinal changes during carcinogenesis, as well as careful consideration of dosing, timing, and potential off-target effects.

In summary, our findings reveal that CINI in oral squamous cell carcinoma establishes maladaptive peripheral nerve crosstalk characterized by sympathetic plasticity, localized NE release, and the acquisition of α1-ADR sensitivity in sensory neurons, which amplifies nociceptive signaling. Importantly, targeted manipulation of sympathetic activity or sensory input disrupted this feed-forward loop, reducing both tumor growth and pain behaviors. Given that several FDA-approved α1-ADR antagonists, including prazosin and doxazosin used in this study, are already in clinical use for hypertension and benign prostatic hyperplasia (BPH) (*118*), these agents may represent promising therapeutic candidates to simultaneously attenuate tumor progression and alleviate cancer-associated pain.

## 4.0 Materials and Methods

### 4.1 Study Design

The aim of the study was to identify if sympathetic-to-sensory neural axis signaling contributes to spontaneous pain behavior in OSCC. Patient data was acquired from patients with SCC in the oral cavity (n=88) or oropharyngeal and neck region (n=8) prior to treatment. We conducted Questionnaire on all 96 patients. Blood samples for NE Elisa were obtained from 56 patients to determine circulating catecholamine levels. IHC on resected tumor samples were acquired from a total of 13 patients with oral cavity tumors and applied to 1 slide/antibody/patient to determine the nerve subtypes innervating the tumors. Resected tongue tumor tissue was also used to determine PNI by a pathologist in 96 patients.

*In vivo* studies in syngeneic orthotopic mouse models and post tissue processing were used to assess molecular mechanism for sympathetic-to-sensory neural axis signaling. Mice groups for all experiments were randomized before inoculation or treatment. Group size was based on prior data to ensure 80% power at α = 0.05. Sample sizes are provided in the figure legends. For human samples, n refers to biologically independent patient samples. For animal experiments, n refers to the number of biologically independent animals per group. For mouse studies, data collection would only be stopped early if tumor burden exceeded over 250mm^3^ in volume. Z-score was determined for potential outliers, with criteria for exclusion being ±3 standard deviation from the mean. There were no reported outliers in the data presented. Endpoints for *in vivo* studies were selected prospectively, with PID12 and PID40 for MOC2 and MOC1 respectively chosen as endpoints due to the similar tumor size growth for comparison and below the threshold for tumor burden cutoff to minimize disease burden. Quantification of immunohistochemical results was performed by an investigator blinded to the groups, using Image J software (NIH) unless otherwise noted in methods section. All key findings were confirmed in at least two independent experiments. Antibodies applied in flow cytometry, IHC, and Western Blot are found in the methods section, while the Taqman Assays are summarized in **Table S4**.

### 4.2. Patient data collection

Prospectively collected patient-reported pain data and blood samples were obtained from 96 oral SCC patients for these analyses; all were >18 years old and underwent surgical resection of the oral cavity. Patients with recurrent disease or second primary cancers were excluded. Demographics and clinical characteristics (**Table 1**) were obtained from medical record review at the pre-treatment/diagnosis timepoint and included: age at diagnosis, sex, race, history of tobacco use, history of alcohol use, tumor site, primary tumor stage, nodal status, extracapsular spread (ECS), PNI (yes/no), and opioid use. The total sample size of 96 oral cavity HNSCC patients was determined to focus on the primary study aim for the targeted domains of pain and psychological symptom burden.

#### 4.2.1. Questionnaires

Patient-reported pain was measured via the Brief Pain Inventory (BPI) and the University of California San Francisco Oral Cancer Pain Questionnaire (UCSF OCPQ). Validation of the OCPQ to assess function-evoked and spontaneous pain has been previously confirmed(*28*). Five subscales of pain were derived from these questionnaires: BPI Mean Pain Severity, BPI Mean Pain Interference, OCPQ Composite Score, OCPQ Function-evoked Pain, and OCPQ Spontaneous Pain. Function-evoked pain was calculated through the sum of questions 1, 3 and 5 and spontaneous pain was calculated through the sum of questions 2,4 and 6; higher score indicates greater symptom severity. Symptoms of depression and anxiety were measured using the Patient Health Questionnaire 8 (PHQ-8) and General Anxiety Disorder 7 (GAD-7), respectively. Higher scores on the PHQ-8 and GAD-7 indicate greater symptom severity. All questionnaires were administered at a preoperative clinic visit prior to analgesia or other treatment.

#### 4.2.2. Blood Collection

Blood from 58 oral SCC patients (pre-operative clinic visit) and 15 healthy volunteers was collected into a lavender top tube, centrifuged at 200 x g for 10 min to isolate plasma, and stored in 1.5mL Eppendorf tubes at -80°C until needed. All patients provided informed written consent; this study was approved by the Institutional Review Board (IRB) associated with the University of Pittsburgh Cancer Institute (STUDY20010205).

#### 4.2.3. Statistical Analysis

To estimate the number of patients needed for this study, we used OCPQ data from 176 HNSCC patients (unpublished) as a key primary endpoint; from the composite score distribution (range 1 – 768), 28% were categorized as low (score ≤ 50). We also use plasma NE concentration; NE concentration in leukoplakia and oral cavity HNSCC patients was 52 ± 6 pg/mL and 462 ± 48 pg/mL respectively (*19*). To test the association of pain with NE, 88 patients are required for 80% power for a two-sample Wilcoxon rank-sum test (equivalent to the Mann-Whitney U test) at α = 0.01. NE is one of several endpoints for the study therefore, the reduced alpha level of .01 is used to reduce overall type I error when analyzing the multiple endpoints proposed for this study. While we are adequately powered to evaluate patient reported pain and psychological symptom burden in HNSCC patients, blood sample acquisition and NE platelet measurement failed to meet 80% power to assess the association of pain with NE (n=58). We preceded with the evaluation and restricted statistical conclusion based on limited sample size. Analysis was performed using R Statistical Software (version 4.1.2; Foundation for Statistical Computing, Vienna, Austria) and RStudio (1.1.456; RStudio, Inc, Boston, Massachusetts). For descriptive statistics, we calculated frequency and percentage for categorical variables and median (Q1, Q3) for continuous variables. Statistical analyses were conducted as regression analyses. We estimated and tested the effect of each pain measure on the log transformed Norepinephrine levels (NE) in linear regression models, after controlling for sex, age, t-stage, and PNI status (dichotomized as presence/absence). We then analyzed the effects of predictors, including depression, anxiety, NE, sex, age, t-stage, n-stage, and ECS, on PNI status in univariable logistic regression models. We also studied the effects of the pain measures on PNI status in multivariable logistic regression models while controlling for NE, t-stage, n-stage, and sex. All hypothesis tests were conducted at the α = 0.05 level and p-values were adjusted using Holm’s method.

### 4.3 Norepinephrine (NE) enzyme-linked immunosorbent assay (ELISA)

To quantify circulating NE in patients with HNSCC, we used platelets isolated from whole blood collected a diagnosis/pre-treatment. Platelets possess the ability to sequester circulating norepinephrine for a long period of time, acting as a biological reservoir and snapshot of overall sympathetic nervous system activity (*29, 30, 119*). *Platelet isolation and lysis:* Whole platelet isolation and lysis were performed at 4°C. Plasma samples were thawed on ice and centrifuged at 1900 x g for 10 minutes to isolate platelets. Red blood cell lysis buffer (Sigma-Aldrich) was applied to the platelet pellets followed by additional centrifugation to remove all blood contamination in pellets. Pellets were stored at -80 °C until needed. Platelet lysis buffer (50µl/sample) containing 2% NP40, 30mM HEPEs, 30mM NaCl, and 2mM EDTA at pH 7.4 supplemented with complete protease inhibitor (Sigma-Aldrich) was subsequently added to platelet pellets and vortexed until no significant pellet was visible. Lysed platelet solution was centrifuged (1900 x g for 5 minutes at 4°C) to isolate any remaining debris and the supernatant was removed and stored on ice for immediate use. *ELISA*: Lysed platelet supernatant solution was diluted 1:2 in PBS^+/+^ for optimal NE preservation. The diluted samples were used in the Norepinephrine Research ELISA kit (Rocky Mountain Diagnostics) and run according to manufacturer’s instructions. Results were normalized to total protein in solution as measured using Piece Rapid Gold BCA Protein Assay Kit (Thermo Fisher Scientific) according to manufacturer’s instructions.

### 4.4 Immunohistochemistry on human tissue

Paraffin-embedded tumor tissue blocks from HNSCC patients were obtained from the Head and Neck Tissue Bank, and sectioned (5um) and stained by the Developmental Pathology Laboratory in the Department of Pathology. Antigen retrieval was performed using a citrate buffer (Dako, Carpinteria, CA). The S100 antibody (rabbit polyclonal, Cat# IR50461–2 DAKO (Agilent), Carpenteria, CA) was applied using a 1:200 dilution at room temperature. The secondary antibody consisted of SignalStain Boost (HRP, Rabbit) (Cell Signaling, Danvers, MA). The substrate used was 3,3, Diaminobenzidine + (Dako). Lastly, the slides were counterstained with Hematoxylin (Dako). The sections were scanned at 10x, 20x, and 40x magnification and S100 labeling was used to identify all nerve bundles. Total S100 immunoreactive area was quantified using Aperio ImageScope software (Leica Biosystems). Within each nerve bundle, CGRP or TH immunoreactive area was quantified and summed for total CGRP or TH area, and the percent of CGRP+ or TH+ area relative to total S100 area was then calculated.

### 4.5 Cell culture

Mouse oral squamous cell carcinoma lines 1 and 2 (MOC1 and MOC2, Kerafast) were cultured in IMDM/F12 (2:1; Gibco) supplemented with 5% Fetal Bovine Serum (Corning), penicillin streptomycin solution (Corning), 5 µg/mL insulin (Sigma-Aldrich), 40 ng/mL hydrocortisone (Sigma-Aldrich), and 5 ng/mL epidermal growth factor (EMD Millipore). Cell lines were cultured in 10 cm diameter tissue-culture treated dishes at 37°C with 5% CO_2_ and passaged using 0.25% trypsin (Corning). Cells were collected for inoculation from a passage number less than 19. Normal cell comparators primary epidermal keratinocytes (Cell Biologics) from C57 mice were cultured in Complete Epithelial Cell Medium Kit including base media and 0.5mL epidermal growth factor, 0.5mL hydrocortisone, 1% antibiotic-antimycotic, 2% FBS (Cell Biologics). Cells were passaged and maintained according to vendor’s instructions, using Gelatin-Based Coating and Trypsin-EDTA 0.05% Solution (Cell Biologics) at 37°C with 5% CO_2._

### 4.6 Animals

Adult (6-12 weeks, 20-30g) male and female C57BL/6 (Strain #000664; Jackson Labs, Bar Harbor, ME) for the majority of the experiments except male and female *Phox2b*-cre:: tdTomato mice (Strain #016223; Jackson Labs, Bar Harbor, ME) and *Npy*-hrGFP mice (Strain #006417; Jackson Labs, Bar Harbor, ME) which were used to assess sympathetic sprouting in to the trigeminal ganglia. All mice were housed in a temperature-controlled room on a 12:12-hour light cycle (0700-1900 hours light), with unrestricted access to food and water. The ARRIVE Essential 10 (*120*) were followed for all preclinical experimental design. Researchers were trained under the Animal Welfare Assurance Program. All procedures were approved by the University of Pittsburgh Institutional Animal Care and Use Committee (IACUC, #24065005) and performed in accordance with the National Institutes of Health Guide for the Care and Use of Laboratory Animals.

### 4.7 Retrograde labeling of tongue primary afferent neurons

The retrograde tracer 1,10-dioctadecyl-3,3,30,30-tetramethy lindocarbocyanine perchlorate (DiI, Invitrogen) or Fast Blue (Polysciences) was injected peripherally into adult C57Bl/6 mice under 3-5% isoflurane anesthesia (Covetrus) within the anterior lateral tongue to retrogradely label tongue afferents. DiI, used for identifying tongue afferents in Ca^2+^ imaging and single cell PCR, was dissolved at 170 mg/mL in DMSO (Thermo Fisher Scientific), diluted 1:10 in 0.9% sterile saline (Hospira) and injected bilaterally using a 30 g needle for a total volume of 5-7µL per tongue. Fast Blue, used for immunofluorescence, was dissolved at 2.5% in sterile dH_2_0 and injected bilaterally for a total volume of 3-4µL per tongue. Retrograde labeling occurred 7-10 days prior to inoculation.

### 4.8 Mouse Model and Manipulations

#### 4.8.1 Orthotopic oral cancer mouse model

To generate the syngeneic orthotopic oral cancer mouse model, adult male and female mice under 3-5% isoflurane anesthesia were inoculated into the anterior lateral portion of the tongue with either 1×10^6^ MOC1 cells or 2×10^4^ MOC2 cells in 30 μL of a mixture of DMEM (Gibco) and Matrigel (Corning) at a 1:1 ratio as previously described (*121*). Injection of DMEM and Matrigel alone was used as a control (i.e. sham). Tumor cells grow and tumor burden is focal to the anterior portion of the tongue. This technique allows for easy access to the entire tumor and clear caliper measurement throughout progression. Tongue tumor dimensions (length, width, and height) were measured using calipers and the volume of the tumors was calculated using the following formula: V=4/3π((length x width x height)/2). The primary limitation of this technique is that animals are repeatedly anesthetized for clinical tongue assessment. However, we ensure that sham mice are also anesthetized in the same manner to control for any off-target impact of anesthesia on tumor growth, immune response, and behavior. Harvest was performed on post inoculation day 14 (PID14) for MOC2 and PID40 for MOC1; mice under 3-5% isoflurane anesthesia were transcardially perfused and processed for subsequent experiments.

#### 4.8.2 Local neuron ablation

Mice were subjected to a double tongue injection protocol using either resiniferatoxin (RTX) or 6-hydroxydopamine (6OHDA) to ablate either sensory or sympathetic neurons respectively. RTX was prepared from a stock solution in vehicle (0.9% NaCl, 0.25% Tween-80, 2mM ascorbic acid) of 100 ng/µl. For each injection, the stock solution was diluted to 50 ng in 30 µl of vehicle. 6OHDA, stored as powder, was prepared fresh for each use by dissolving 1 mg in 160 µl of 1% ascorbic acid solution, yielding a dose of 6 mg/kg for a 20 g mouse. The solution was protected from light during preparation and administration. For both toxins, two injections were performed: an anterior tongue injection on Day 1, followed by a posterior tongue injection on Day 3, with a recovery day in between to minimize inflammation. Total volume for each solution was 30uL delivered with an insulin syringe. Following the Day 3 injection, mice were allowed to recover for one week prior to retrograde labeling, followed by an additional week of recovery before inoculation.

#### 4.8.3 Surgical Sympathectomy

Mice were anesthetized with an intraperitoneal (i.p.) injection of ketamine/xylazine (80 mg/kg and 10 mg/kg, respectively) and positioned supine on a dissection mat. The ventral surface of the neck was shaved and thoroughly cleaned with 70% ethanol. Ophthalmic ointment was applied to prevent corneal drying. A ventral midline incision was made on the neck using a sterile scalpel, and underlying fascia, submandibular salivary glands and hyoid muscles were retracted. and the surgical field was stabilized open. Blunt dissection with forceps and spring scissors was used to access the SCG, located medial to the common carotid bifurcation. Using fine forceps and spring microscissors, the SCG was resected without vascular injury. The procedure was repeated bilaterally. After resection, the incision was closed with absorbable sutures. Triple antibiotic ointment (TAO) was applied topically to the incision site, and 500 μL of sterile saline was administered subcutaneously at the nape of the neck for postoperative hydration. Animals received i.p. 0.05mg/kg buprenorphine for analgesia once daily for 3 days. Following the surgery, mice were allowed to recover for one week prior to retrograde labeling, followed by an additional week of recovery before inoculation.

### 4.9 Mouse Behavior Assays

#### 4.9.1 Grimace

Spontaneous nociceptive behavior was measured using the grimace assay and automated analysis software. Mice were acclimated to experimental room in home cages for 20 minutes before any behavior. Mice were then placed inside small behavioral chambers (Hypothesis to Hardware) that allow free movement and video recorded for 20min (Sony HDR CX405 Handycam); mouse facial features, referred to as action units (eyes, ears, nose, whiskers), associated with the mouse grimace scale are then quantified using the *PainFace* platform (*40*). Mean score and area under the curve (AUC) were obtained for the first 200 usable frames as defined by n=3 or 4 identified action units. Grimacing bouts were defined as the total number of grimacing frames in the first 200 frames, in which the facial action unit score ≥1.5 standard deviations from baseline. The number of animals by sex used was based on pilot data of n=3/sex/group. For sample size and power calculations sham mice exhibited a mean AUC of 312.25 with a standard deviation of 50.05, while MOC2 tumor bearing mice exhibited a mean AUC value of 552.75 with a standard deviation of 157.85 with no significant differences between sexes (p=0.7000). G*Power analysis yielded an effect size of d = 2.0642, requiring n = 4 per group to achieve >80% power at α = 0.05.

#### 4.9.2 Dolognawmeter

The dolognawmeter assay and automated device quantifies gnawing activity. The outcome variable (gnaw-time) is the time required by a rodent to gnaw through the second of two obstructing polymer dowels in series (a discrete gnawing task) that block escape of a rodent confined in a narrow tube. Extended gnaw-time relative to baseline values is a validated index of orofacial nociception in mice with oral cancer (*38*). Each mouse is placed into a confinement tube where forward movement of the rodent in the tube is obstructed by 2 polymer dowels. The mouse voluntarily gnaws through both dowels to escape the device. The second of two sequentially obstructing polymer dowels is connected to a digital timer. The timer automatically records the duration required for the mouse to sever the second dowel. To acclimatize the mice and improve consistency in gnawing behavior, all mice were trained for 7-9 sessions in the dolognawmeter. Training was accomplished by placing the mice in the device and allowing them to gnaw through the obstructing dowels in the same manner as the subsequent experimental gnawing trials. For the orthotopic oral cancer pain model, a baseline gnaw-time (mean of the final 3 training sessions) was established for each mouse followed by behavioral testing twice per week for up to 6 weeks. Sample size and power calculations were based on historical data from previous publications (*21, 122*) as this behavior is used routinely in the lab. A change greater than 50% of baseline will be considered a significant difference in pain behavior. From historical data sham female mice exhibited a mean percent change in gnaw-time of 0.59 with a standard deviation of 16.66, while MOC1 tumor bearing female mice at PID40 exhibited a mean percent change in gnaw-time of 318.0 with a standard deviation of 247.7. G*Power analysis yielded an effect size of d = 1.808, requiring n = 5 per group to achieve >80% power at α = 0.05. Each mouse was compared to its own baseline gnaw-time, and data are presented as a percent change ± standard error of the mean.

#### 4.9.3 Open field test

Open field testing was conducted using two open field chambers and ANY-maze behavioral tracking software (Stoelting Co.). Mice were tested in a darkened box mounted on metal bases, with overhead cameras connected to a USB hub and behavior computer. Before each session, a background image was captured, followed by a 1-minute acclimation and a 15-minute test period. After testing, videos were analyzed by ANY-maze; results captured included: distance traveled, average speed, freezing episodes and duration, immobility, and time spent in 3 zones (middle, edge, center). The number of animals by sex used for OFT freezing behavior was based on pilot experiments using MOC2 tumor bearing mice (n=2-5/sex/treatment group). For the sample size and power calculations the males had a mean 32 freezing bouts with a standard deviation of 3.6, while female mice had a mean 39.33 freezing bouts with a standard deviation of 3.06 and no significant difference between groups (p=0.1000). Using G*Power software, this yielded a robust effect size of d = 2.19. A sample size of n = 5 per group was calculated to achieve >80% power at α = 0.05. Track plots and heatmaps were generated by the ANY-maze software.

### 4.10 Immunohistochemistry (IHC)

At least 7 days after retrograde labeling, mice were perfused with PBS (Gibco) followed by 4% paraformaldehyde (PFA; Electron Microscopy Sciences). Tongue tissue, bilateral trigeminal ganglia (*93*), and superior cervical ganglia (SCG) were removed.

#### 4.10.1 Ganglia tissue – slide mounted

Ganglia were post-fixed for 1 hour in 4% PFA, cryoprotected in 30% sucrose at 4°C overnight, embedded in OCT compound (Tissue-Tek cryosectioned (10-12 µm), and mounted on Superfrost Plus slides (Thermo Fisher Scientific). To identify injured neurons or sympathetic sprouting, slides were subsequently incubated in blocking solution comprised of PBS containing 10% normal goat serum (Gibco), 0.01% Triton-X 100 (Sigma-Aldrich), 1% bovine serum albumin (Fisher Scientific), and 0.1% Tween-20 (Sigma-Aldrich) for 1 hour at room temperature (RT) followed by incubation overnight in either rabbit anti-ATF3 (1:250, Abcam) or rabbit anti-tyrosine hydroxylase (TH, 1:500, Millipore Sigma). Following primary antibody, slides were extensively washed in PBS and incubated in goat anti-rabbit Alexa Fluor 594 (1:250, Jackson ImmunoResearch) for 2 hours at RT and cover-slipped with Fluoro-Gel mounting media (Electron Microscopy Sciences). Using a Keyence BZ-X810 microscope with Keyence Imaging software, TG and SCG sections were imaged at either 10x or 20x magnification; TG sections were imaged within the intersection of the mandibular and maxillary branch where most retrograde labeled trigeminal tongue neurons reside. Ganglion neurons with distinct nuclei and at least 50% of the cell area labeled with DiI or Fast Blue were counted in every fifth section (9 sections/mouse/treatment group). ImageJ software (NIH) was used to count retrograde labeled neurons that overlapped with protein-specific immunoreactivity per animal. ImageJ software and the NeuronJ plugin were used to trace and measure total sympathetic sprouting.

#### 4.10.2 Tongue tissue – slide mounted

Tongue tissue was cryoprotected similarly to ganglia tissue and sectioned at 20µm thickness. Primary antibodies used to assess sympathetic and sensory innervation were sheep anti-TH (1:250, Millipore Sigma), or rabbit anti-CGRP (1:250, Cell Signaling Technology) respectively; rabbit anti-PGP9.5 (Boster Bio) was used as a pan-neuronal stain and mouse anti-GFAP (1:250, Sigma-Aldrich) was used to identify an injury response. Incubation in goat anti-sheep Alexa Fluor 647 and goat anti-rabbit Alexa Fluor 800 (1:250, at 4°C for 24 hours) were used for visualization. GFAP signal intensity quantification was performed using the open-source software Fiji (ImageJ). Images were first converted to a 16-bit format to preserve the dynamic range of data. All images were acquired using identical exposure settings and processed using a standardized workflow. For each image, Regions of Interest (ROIs) were defined using the ROI Manager and manually traced to not include the border to avoid edge effects being quantified. To measure intensity, the Set Measurements function was configured to capture only the Mean Gray Value. This value represents the average intensity of all pixels within the defined ROI, calculated as: Mean Gray Value= Σpixel intensities/total number of pixels in ROI.

#### 4.10.3 Tongue tissue – optically cleared

Samples were optically cleared according to previously published protocols using CUBIC chemistry; all solutions were prepared according to recommendations(*123*). Cleared tongues were embedded in 6% agarose gel and sectioned on a sagittal or coronal plane at 300µm thickness using on a vibratome. Sections were then incubated in CUBIC R1 solution for 48-72 hours followed by staining to identify all peripheral nerves as well as specifically sympathetic or sensory innervation. Due to high autofluorescence of tongue muscle, antibody multiplexing was not possible. For antibody staining, free floating tongue sections were incubated in blocking solution containing 6% normal goat serum and 1% Triton X-100 overnight at RT followed by incubation in one of the following primary antibodies for 72 hours at 4°C: rabbit anti-PGP9.5 (1:250, Boster Bio), rabbit anti-CGRP (1:250, Cell Signaling Technology), rabbit anti-TH (1:250, Millipore Sigma). Samples were incubated in secondary antibody goat anti-rabbit Alexa Fluor 647 (1:250, Jackson ImmunoResearch) at 4°C for 24 hours. The entire thickness of each sample was imaged using the Z-stack function on the Keyence BZ-X810 microscope; stacked images were subsequently stitched and full-focus processed using Keyence Imaging software.

### 4.11 Western blot

Western blot was performed using tongue tissue from sham or MOC2 inoculated male and female mice. Samples were homogenized in RIPA Buffer with 1% Halt Protease Inhibitor Cocktail (Thermo Scientific) and lysed for 2 hours at 4C; supernatant was recovered after centrifugation at 16,000 x g for 10 min and frozen at -80C until use. Protein concentration was measured prior to Western blotting via Pierce Rapid Gold BCA Protein Assay (Thermo Scientific). Samples were loaded at 50ug/well into Bolt Bis-tris Plus 4-12% Mini Protein Gels (Invitrogen) and transferred to PVDF using iBlot 2 semi-dry transfer system (Invitrogen) according to manufacturer’s instructions. Membrane was blocked for 1 hour using Fluorescent Blocking Buffer (Thermo Scientific) then co-incubated with primary antibodies (mouse anti-GFAP, 1:1000 (Sigma-Aldrich); rabbit anti-GAPDH, 1:50,000 (ThermoFisher, loading control)) overnight at 4C. Membrane was washed and incubated for 1 hour with secondary antibodies (goat anti-Mouse Alexa Fluor 680, 1:10,000 (ThermoFisher); goat anti-rabbit DyLight 800, 1:10,000 (ThermoFisher)). Membrane was washed and transferred to 1xTBS for imaging using Odyssey CLx (Li-Cor). Gel analysis was performed using ImageJ (NIH), using GAPDH as the loading control protein across groups.

### 4.12 Trigeminal ganglion and superior cervical ganglion dissociation

TG dissociation was used to generate neurons for single cell pick up for PCR, Ca^2+^ imaging and quantification of TG infiltrating immune cells by flow cytometry. SCG dissociation was used to generate neurons for Ca^2+^ imaging. Mice under 3-5% isoflurane anesthesia were transcardially perfused with Hank’s Balanced Salt Solution (Gibco) containing 5mM HEPES (HBSS+H, Thermo Fisher Scientific). TG tissue was dissected and dissociated for Ca^2+^ imaging as previously described(*124*). Briefly, tissue was harvested into HBSS+H, minced, and subjected to a two-step enzymatic digestion consisting of L-cysteine and papain (Sigma-Aldrich) for 10 minutes at 37°C followed by a combined collagenase type II and dispase II (Gibco) digestion for 20 minutes at 37°C. Tissue was then centrifuged at 240xg and passed through a Percoll gradient (1000xg centrifugation, GE Healthcare) to isolate neurons, resuspended in DMEM/F12 (Gibco), and plated on 12mm poly-L-ornithine and laminin (Sigma-Aldrich) coated glass coverslips (Electron Microscopy Sciences). SCG tissue was dissected and dissociated for Ca^2+^ imaging as follows: SCG tissue was harvested into HBSS+H, minced and subjected to a two-step enzymatic digestion consisting of L-cysteine and papain for 10 minutes at 37°C followed by a combined collagenase type II and dispase II digestion for 20 minutes at 37°C. Tissue was then centrifuged at 240xg and passed through two fire polished pipettes of decreasing size to isolate neurons, resuspended in DMEM/F12, and plated on 12mm poly-L-ornithine and laminin coated glass coverslips.

### 4.13 Quantitative PCR (qPCR)

*For single cell analysis*, dissociated neurons were collected within 1 h of removal from the incubator and within 8 h of removal from the animals. DiI-positive single neurons were identified using fluorescence microscopy, picked up using glass capillaries (World Precision Instruments) held by micromanipulator (Sutter Instruments) using headstage (EPC10 HEKA) and electrode holder under brightfield optics. Cell size selection criterion was ≤30µm. Each cell was transferred into a 0.2 ml PCR tube containing 9μl of single cell lysis solution and DNase I from Single Cell-to-CT^TM^ Kit (Invitrogen). After 5-minute incubation, 1μl of stop solution was added, cells were incubated for 2 minutes, then immediately snap frozen and stored at -80°C until further use. For *in vitro* cell line analysis, cells were harvested from a passage less than 18, pelleted, snap frozen, and stored at -80C until use. *For whole ganglia and cell line analysis*, total RNA was isolated from TG or cell pellets using the Qiagen RNeasy Plus Mini Kit (Qiagen Inc.); SCG RNA extractions were performed using the RNA Microprep Plus Kit (Zymo Research) according to manufacturer’s instructions. For all samples, reverse transcription using Quantitect Reverse Transcription Kit (Qiagen Inc.); cDNA pre-amplification (single cell <u>only</u>), and qPCR were executed per manufacturer’s instructions. Relative expression levels of genes of interest were assessed using TaqMan Gene Expression Assays (**Table S4)** and either TaqMan Gene Expression Master Mix (Thermo Fisher Scientific) for single cell or Taqman Fast Advanced Master Mix (Thermo Fisher Scientific) for whole ganglia, using a 96 well Quantstudio 3 Real-Time PCR System (Thermo Fisher Scientific). *Gapdh* or *GusB* were used as internal control genes. Any cell with a *Gapdh* cycle threshold (Ct) of 27 or higher was excluded from further analysis. For single cell analysis, genes were considered ‘not expressed’ if one sample either failed to detect expression or the Ct was above 35. All samples were run in duplicate or triplicate to account for pipetting errors. Relative fold change of gene expression data in cancer mice compared to sham mice was calculated using the 2^−ΔΔCt^ method.

### 4.14 Analytical flow cytometry

Analytical flow cytometry was used to assess macrophage infiltration into TG tissue from sham and MOC2-bearing mice. Immune cells dissociated from TG were first incubated for 20 minutes at RT in the dark with Zombie NIR viability label (1:1000, Thermo Fisher Scientific) followed by a wash with staining media [PBS (Gibco) containing 3% FBS (Corning), 1mM EDTA (Sigma)]. To stain extracellular markers, cells were incubated at 4°C in the dark for 30 minutes with the following antibodies diluted 1:100 in staining media: CD45 BUV-395 (BD Biosciences), CD11b APC(Biolegend), and F4/80-BV 785 (Biolegend), followed by a wash with staining media. For intracellular staining, cells were then incubated at 4°C in the dark for 30 minutes in fix/permeabilization buffer (Miltenyi Biotec), then blocked with 5% normal mouse serum (Invitrogen) in 1x permeabilization buffer (Miltenyi Biotec) for 10 minutes at RT in the dark. CD206 PE (Phycoerythrin) /Cy7 antibody (1:50, BioLegend) was added to cells for 30 minutes at 4 °C in the dark. Leukocytes from the spleen were used for compensation controls. An average of 518,032 ± 33,483.79 live cells per sample were acquired on a 5-laser Becton Dickenson LSR Fortessa II analyzer (BD Biosciences) and data were analyzed with FlowJo™ (v10, Tree Star).

### 4.15 Calcium imaging

TG neurons were dissociated and incubated with 5µM Fura-2AM (Fura, Invitrogen), a Ca^2+^ indicator, at RT for 30 minutes. Coverslips were placed in a Normal Bath (NB, 130mM NaCl, 3mM KCl, 2.5mM Cacl_2_, 0.6mM MgCl_2_, 10mM HEPES, 10mM Glucose, pH7.4, 325mOsm) wash for 5 minutes. Coverslips were then placed in the recording chamber and perfused continuously with NB. Baseline Ca^2+^ and spontaneous activity were measured for one minute prior to any stimulus. Spontaneous activity was defined as a change in Fura (ΔF) ratio (340/380) 20% of baseline occurring 2 or more times in the initial 60-second baseline measurement. To measure vehicle control, 0.001% HCl in NB was applied to the cells for 30 seconds. To measure norepinephrine (NE) response, 10µM NE (Sigma-Aldrich) in NB was applied to the cells for 30 seconds. The number of animals by sex used for the quantification of evoked response was based on pilot data from n=3/sex/group with ≥15 tracer positive neurons assessed per animal. Using G*Power software from pilot data yielded a robust effect size of d = 8.910467. A sample size of n = 4 per group was calculated to achieve >80% power at α = 0.05 . Phenylephrine (α1 agonist, 10µM, Tocris), Clonidine (α2 agonist,10µM, Sigma-Aldrich), Doxazosin mesylate, a high-affinity adrenergic α1 selective antagonist (10µM, Tocris), or Acetylcholine (30uM, Sigma-Aldrich) was added to the normal bath perfusion for adrenergic pharmacology testing. KCl (30mM) in NB was applied for 10 seconds to test cell viability at the end of each experiment. Response to stimulus was defined as a ΔF 20% of baseline. The magnitude of the response was calculated as (peak ΔF – baseline ΔF). Fluorescence data were acquired by Leica DMi8 microscope at 340nm and 380nm excitation wavelength and analyzed with Leica Application Suite software (Leica Microsystems).

### 4.16 Prazosin-Bodipy

TG neurons were dissociated and incubated with 2.5nM Prazosin-Bodipy (Invitrogen), a highly selective alpha-1 antagonist at RT for 45 min. Coverslips were placed in a Normal Bath (NB, 130mM NaCl, 3mM KCl, 2.5mM Cacl_2_, 0.6mM MgCl_2_, 10mM HEPES, 10mM Glucose, pH7.4, 325mOsm) wash for 5 minutes. Using a Keyence BZ-X810 microscope with Keyence Imaging software, TG neurons were imaged at either 10x or 20x magnification.

### 4.17 Norepinephrine (NE) mass spectrometry

*Serum - Sample preparation:* Metabolic quenching and metabolite pool extraction was performed by adding an equal volume of acetonitrile (ACN) to 50µL serum. Deuterated (13C1)-creatinine and (D3)-alanine (Sigma-Aldrich) were added to the sample lysates as an internal standard for a final concentration of 10µM. After 3 minutes of vortexing, the supernatant was cleared of protein by centrifugation at 16,000xg. 3µL of cleared supernatant was subjected to online LC-MS analysis. A calibration curve of NE was prepared in 50% ACN with deuterated internal standards from 100 pmol/µL to 0.6 fmol/µL. *Tumor Tissue - Sample preparation*: Metabolic quenching and metabolite pool extraction was performed by adding 50% acetonitrile (ACN) at a ratio of 1:15 (wt:vol) in MPBiomedicals Matrix A tubes. Deuterated (13C1)-creatinine and (D3)-alanine (Sigma-Aldrich) were added to the sample lysates as an internal standard for a final concentration of 10µM. Samples were homogenized at 60Hz for 40 seconds using a FastPrep 24 (MPBiomedical) homogenizer before the supernatant was cleared of protein by centrifugation at 16,000xg. 3µL of cleared supernatant was subjected to online LC-MS analysis. A calibration curve of NE was prepared in 50% ACN with deuterated internal standards from 100 pmol/µL to 0.6 fmol/µL. *LC-HRMS Method*: Analyses were performed by untargeted LC-HRMS. The number of animals by sex used for the quantification of intratumoral NE by mass spectrometry was based on pilot data of tongue tissue from n=2-3/sex/group. Using G*Power software from pilot data yielded a robust effect size of d = 1.502562. A sample size of n = 9 per group was calculated to achieve >80% power at α = 0.05. Briefly, samples were injected via a Thermo Vanquish UHPLC and separated over a reversed phase Phenomenex Kinetix C18 column (2.1×150mm, 3μm particle size) maintained at 55°C. For the 15-minute LC gradient, the mobile phase consisted of the following: solvent A (water / 0.1% FA) and solvent B (ACN / 0.1% FA). The gradient was the following: 0-2min 1% B, increase to 5%B over 4 minutes, continue increasing to 98%B over 5 minutes, hold at 98%B for 2 minutes, reequillibrate at 1%B for three minutes. The Thermo Exploris 240 mass spectrometer was operated in positive ion mode, scanning in ddMS2 mode (2 μscans) from 10 to 1000 m/z at 120,000 resolution with an AGC target of 2e5 for full scan, 2e4 for ms2 scans using HCD fragmentation at stepped 15,35,50 collision energies. Source ionization setting was 3.0 and 2.4kV spray voltage respectively for positive and negative mode. The source gas parameters were 50 sheath gas, 12 auxiliary gas at 320°C, and 8 sweep gas. Calibration was performed prior to analysis using the PierceTM FlexMix Ion Calibration Solutions (Thermo Fisher Scientific). Integrated peak areas were then extracted manually using Quan Browser (Thermo Fisher Xcalibur ver. 2.7). Extracted peak areas were then normalized to deuterated internals standards before conversion to concentration using the calibration curve.

### 4.18 Statistics

Statistical significance was set at p < 0.05. Preclinical statistical analyses were performed using Prism (version 10.5.0) statistical software (Graphpad Software Inc.). Pilot studies and power calculations were conducted for all major preclinical studies involving groups of mice. Details are provided within the corresponding methods sections. Box/scatter configurations were used to show the biological variability when illustrative. Given the previous literature demonstrating a sex difference in oral cancer-evoked nociceptive behavior (*122, 125, 126*), all studies were powered to detect a sex difference, and a two-way analysis of variance (*123*) was employed to evaluate an interaction between sex and treatment. In the instance that no effect of sex was found, samples were pooled. For parametric data, results were presented as mean ± standard error of the mean. One-way ANOVA or independent student’s t test was employed to evaluate the difference between groups. To adjust for multiple comparisons, the post-hoc Holm-Sidak test statistic was employed. For non-parametric data, results were presented as mean ± 95% confidence interval. Kruskal-Wallis test with Dunn’s multiple comparisons test was used to test statistical differences in catecholamine concentration between oral SCC patients and healthy controls and to test statistical differences in relative gene expression distribution in sensory neurons between treatment groups. Mann Whitney U test was used to test for a difference between retrograde labeled neurons in SCG sections from MOC2 and sham mice.

## List of Supplementary Materials

Supplemental Figure 1: Tongue tumor-induced nociceptive behavior

Supplemental Figure 2: Sensory nerve injury in tumor bearing mice

Supplemental Figure 3: Adrenergic receptor pharmacology and gene expression

Supplemental Figure 4: Sympathectomy decreased nociceptive behaviors

Supplemental Figure 5: Sensory neuron ablation does not affect anxiety-like behaviors or acquired adrenergic sensitivity

Supplemental Table 1: Multivariable Regression: Pain Effects on Log of NE

Supplemental Table 2: Univariable Logistic Regression: Predictor Effects on PNI Status

Supplemental Table 3: Multivariable Logistic Regression: Pain Effects on PNI Status

Supplemental Table 4: Taqman Assays for single cell and whole ganglia quantitative PCR

## Acknowledgements

We thank the Hillman Cancer Center Head and Neck Tissue Registry for supplying clinical specimens as well as Dr. Elizabeth Bilodeau DMD, MD, MSEd for her expertise in pathological assessment in selection of HNSCC tumor tissue.

## Author contributions

- Conceptualization: Nicole N. Scheff, Andre A. Martel-Matos
- Methodology: Nicole N. Scheff, Marci L. Nilsen, Nicole L. Horan, Andre A. Martel-Matos
- Investigation: Andre A. Martel-Matos, Lisa A. McIlvried, Nicole L. Horan, Nicole A. Rodriguez, Jared I. Rothberg, Megan A. Atherton, Marci L. Nilsen, Nicole N. Scheff
- Visualization: Andre A. Martel-Matos, Nicole N. Scheff, Stephen V. Glass
- Funding acquisition: Andre A. Martel-Matos, Nicole N. Scheff
- Project administration: Andre A. Martel-Matos, Nicole N. Scheff, Marci L. Nilsen
- Supervision: Nicole N. Scheff, Marci L. Nilsen
- Writing – original draft: Andre A. Martel-Matos, Nicole N. Scheff
- Writing – review & editing: Andre A. Martel-Matos, Lisa A. McIlvried, Marci L. Nilsen, Nicole N. Scheff,

## Data and materials availability

All data are available in the main text or the supplementary materials.

