## Supplemental Materials for "Cancer-induced Nerve Injury Unveils a Sympathetic-to-Sensory Nerve Axis in Head and Neck Cancer"

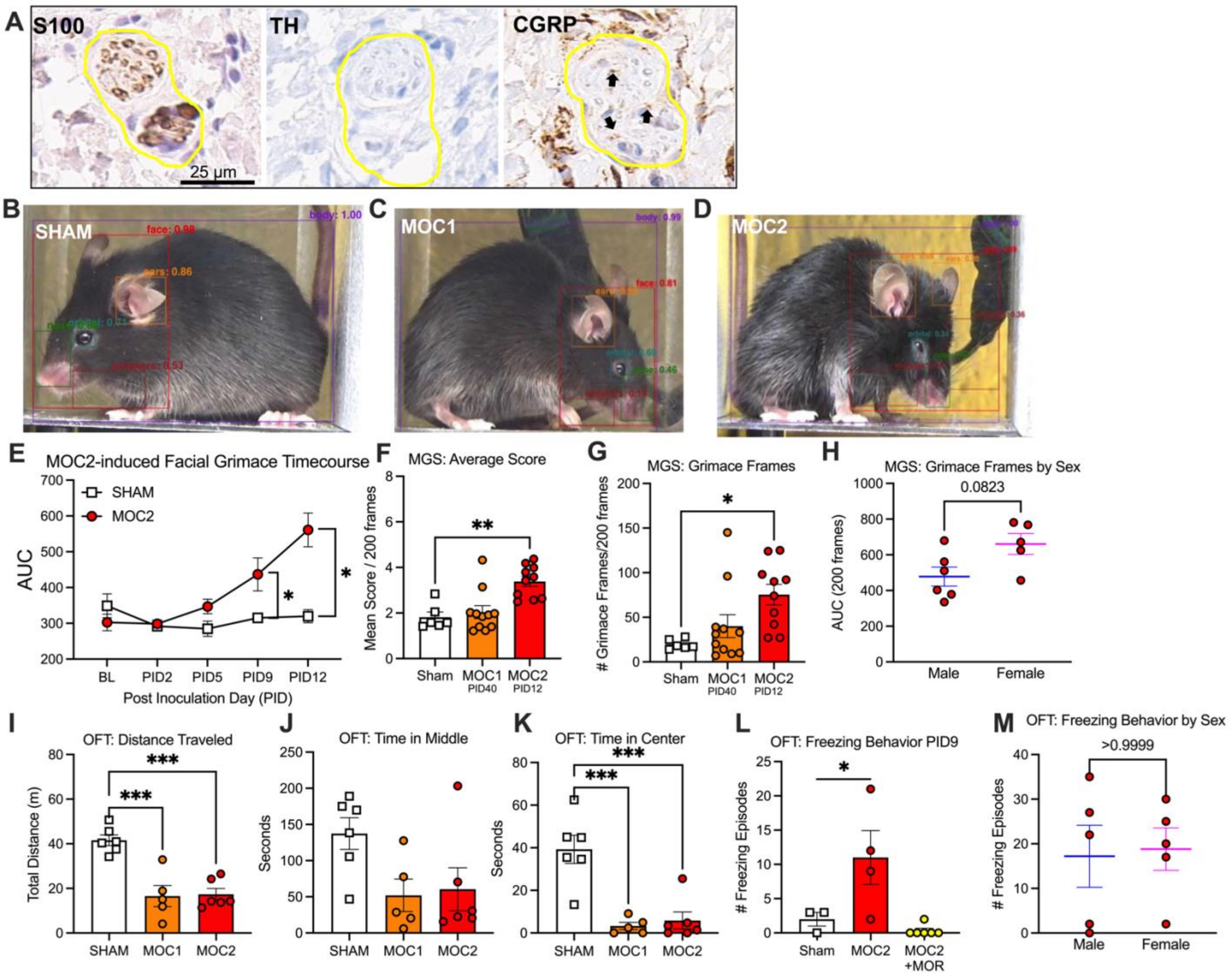


**Supplemental Figure 1: Tongue tumor-induced nociceptive behavior**
(A) Representative image of S100 (left), tyrosine hydroxylase (TH, middle), and CGRP (right) immunoreactivity (IR) in HNSCC tumor section (Male, Stage III). Nerve bundles are outlined in yellow. Positive stain is indicated by black arrows. Image magnification = 20x.

(B-C) Representative video frames from Sham, MOC1 and MOC2 tumor-bearing mouse pulled from recordings for analysis of grimacing behavior, analyzed using PainFace software. The software automatically identified the body and facial region, followed by scoring of individual facial action units (*40*), including ears, orbital tightening (eyes), nose bulge, and whisker position. Sham frame received a cumulative grimace score of 3. MOC1 frame received a cumulative grimace score of 4. MOC2 frame received a cumulative grimace score of 7.

(E) Area under the curve (AUC) analysis of grimace scores over time in MOC2 versus sham mice demonstrates elevated spontaneous nociceptive behavior in MOC2 tumor-bearing mice compared to sham. n=3-5/sex/group, Two-Way ANOVA, Cancer by time interaction, *p<0.05

(F) Mean grimace score over the first 200 usable frames and number of grimacing frames determined as 1.5 times above the standard deviation of the mean baseline grimace score. n=3-5/sex/group, One-Way ANOVA, *p<0.05, **p<0.01.

(G) Number of grimace frames defined as ≥1.5 standard deviations from baseline for sham, MOC1 and MOC2 mice. MOC2 tumor-bearing mice demonstrated a significant increase in the number of grimace frames compared to sham. n=3-5/sex/group, One-Way ANOVA, *p<0.05

(H) Area under the curve (AUC) analysis of grimace scores in MOC2 mice separated by sex demonstrates no significant difference in grimace scores. These data are regraphed from Figure 2G. n=5-6 group, Independent T-test, p>0.05.

(I-K) Open Field Test anxiety-related measures were also analyzed, including total distance traveled (**D**), time spent in the middle zone (**E**), and time spent in the center zone (**F**) of the chamber. Tumor bearing mice display sign of anxiety like behavior, which can be hard to disentangle from nociceptive behavior and tumor burden. n=2-3/group, One-Way ANOVA, ***p<0.005.

(L) Validation of spontaneous freezing behavior in MOC2 tumor-bearing mice PID9, quantified using the Open Field Test and analyzed with ANY-maze software. Freezing episodes of MOC2 tumor bearing mice were significantly reduced by a single IP injection of morphine (10 mg/kg) administered 45 minutes prior to testing, confirming the nociceptive nature of this behavior. n=1-3/sex/group, One-Way ANOVA, *p<0.05

(M) Freezing episodes from MOC2 tumor bearing mice between sexes demonstrate no significant difference. These data are regraphed from Figure 2I. n=5 group, Independent T-test, p>0.05.

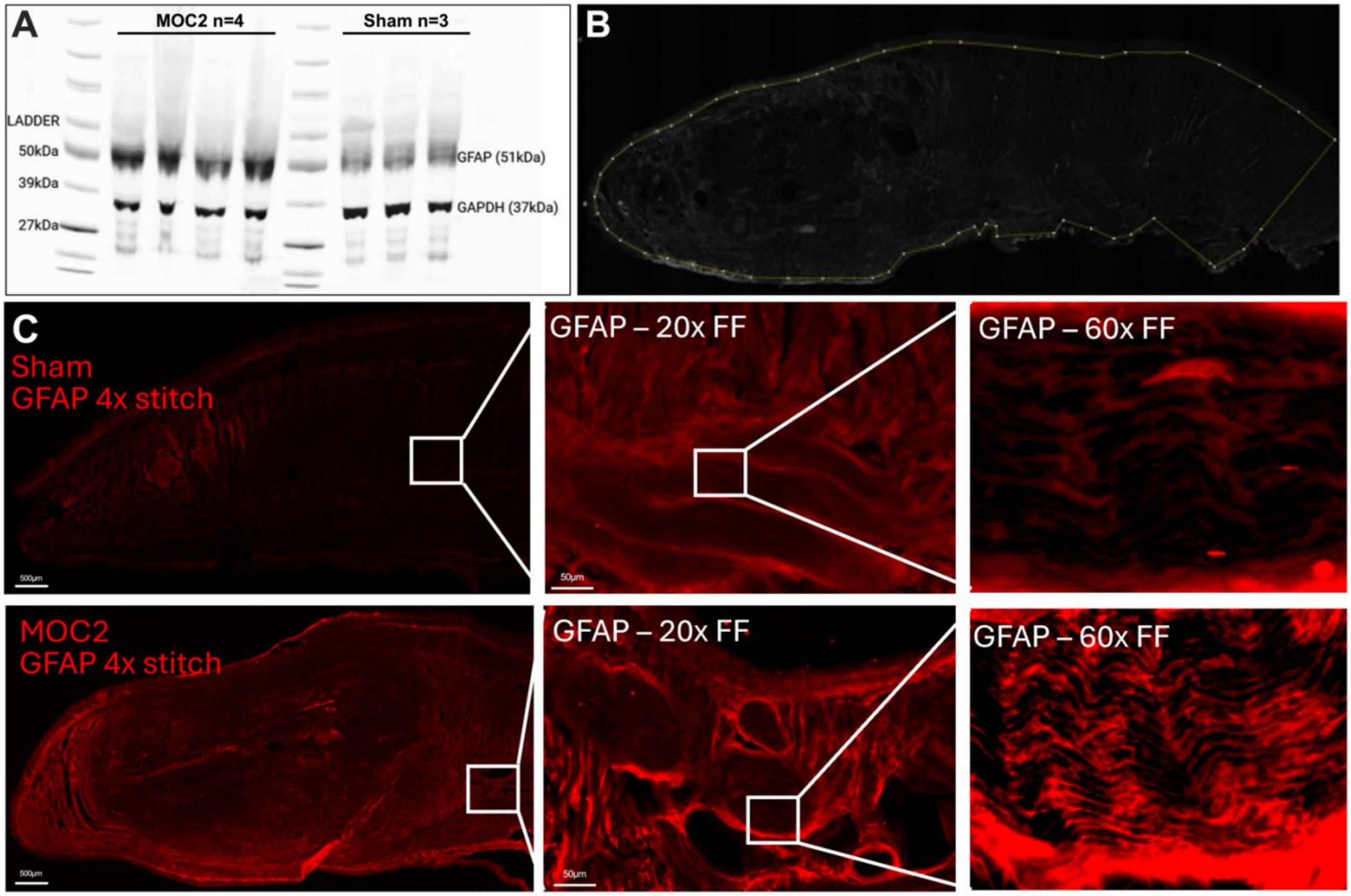


**Supplemental Figure 2: GFAP protein quantification and immunofluorescence at varying magnification**
(A) Full western blot image of GFAP-immunoreactivity in tongue lysates from MOC2 tumor-bearing and sham mice. GAPDH was used as a loading control and is indicated on the blot. Molecular weight markers are indicated on ladder in the left-most lane. MOC2 (n=4 tongue samples); Sham (n=3 tongue samples).

(B) Representative image from quantification analysis of GFAP staining using 20μm sagittal sham tongue section stained with anti-GFAP. Region of interest (ROI) that excludes the edge of the tissue were manually drawn to minimize edge effect impact on mean grey intensity analysis.

(C) Immunohistochemical staining of GFAP-immunoreactivity in 20μm tongue tissue from sham and MOC2 tumor-bearing mice captured at a full section stitch at 4x magnification, a 20x full focus z-stack to account for tissue thickness, and a 60x full focus z-stack to account for tissue thickness. While boxes indicate roughly at what location within the tissue the increased magnification image was taken. The tissues were sectioned and stained within the same slide. Exposure was kept consistent between tissues.

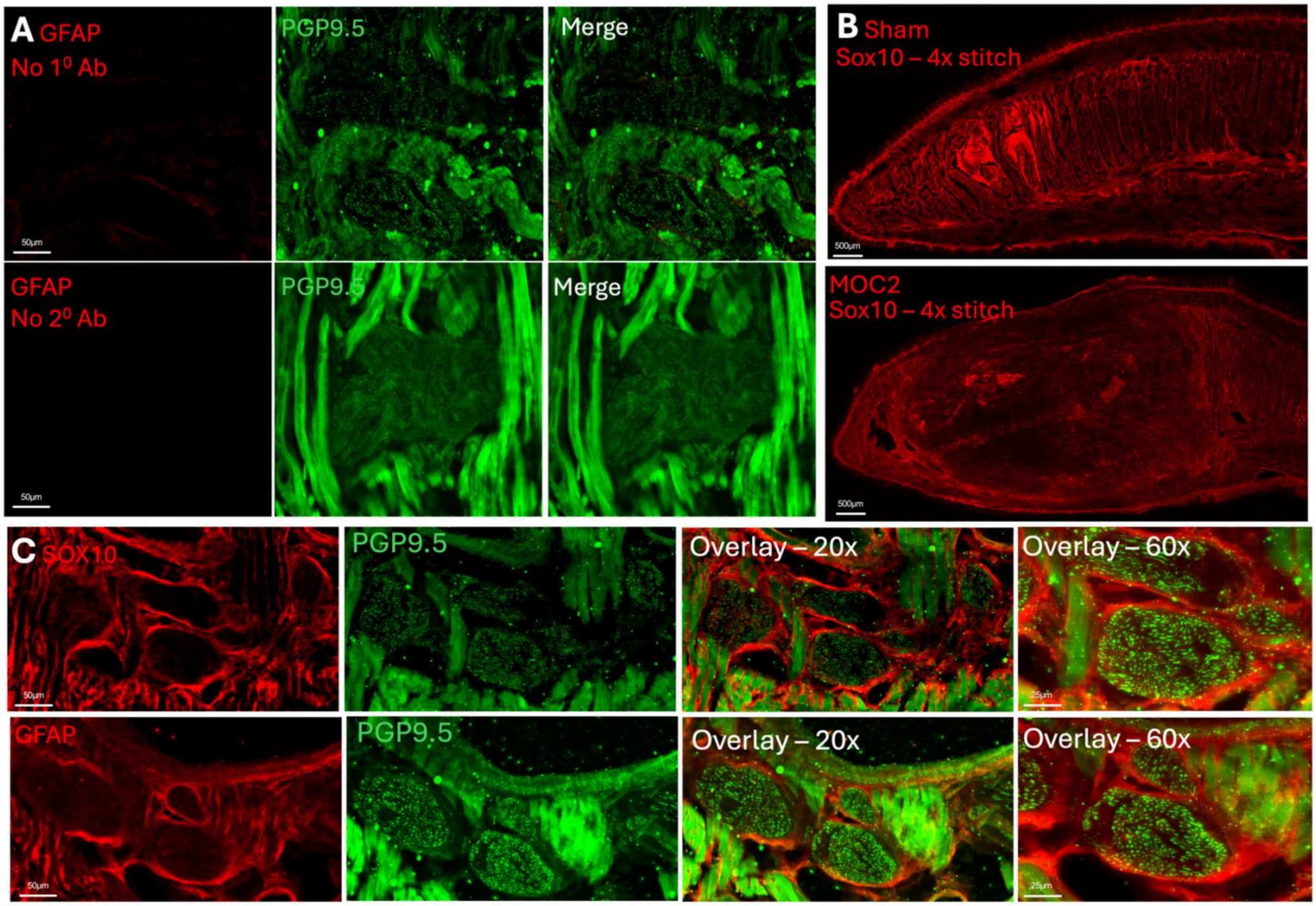


**Supplemental Figure 3: Positive control experiments for GFAP immunofluorescence.**
(A) Antibody controls for GFAP-immunoreactivity were completed in combination with pan-neuronal antibody PGP9.5 to confirm lack of staining around nerve morphology.

(B) Pan-Schwann cell marker, SOX10, was used to label all Schwann cells within the tongue tissue regardless of activation status in sham and MOC2 tongue tumor tissue. Images were taken as a full tissue section stitch at 4x magnification.

(C) SOX10 and GFAP were co-stained with PGP9.5 in serial tongue sections within MOC2 tumor tissue to demonstrate similar immunoreactivity with both antibodies around nerve bundles.

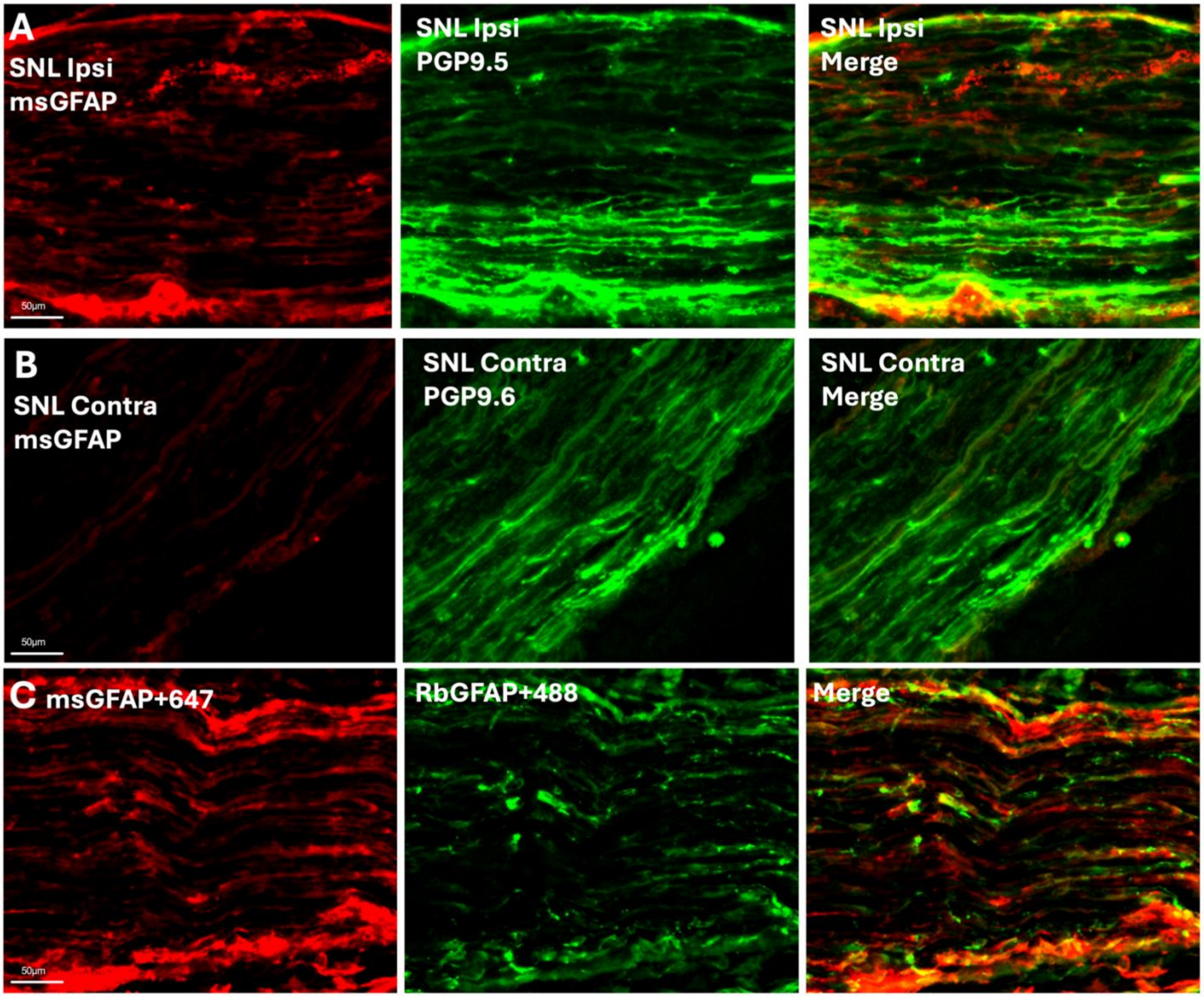


**Supplemental Figure 4: GFAP immunofluorescence in ipsilateral and contralateral sciatic nerve following sciatic nerve ligation (SNL).**

(A) Ipsilateral sciatic nerve section (12µm) stained with co-stained with GFAP and PGP9.5 reveals robust GFAP immunofluorescence on the outside and within the nerve.

(B) Contralateral sciatic nerve section (12µm) stained with co-stained with GFAP and PGP9.5 has minimal GFAP immunofluorescence within the nerve.

(C) Co-staining of two GFAP antibodies raised in two different species from two different suppliers demonstrating similar immunofluorescent staining patterns within the ipsilateral sciatic nerve following SNL.

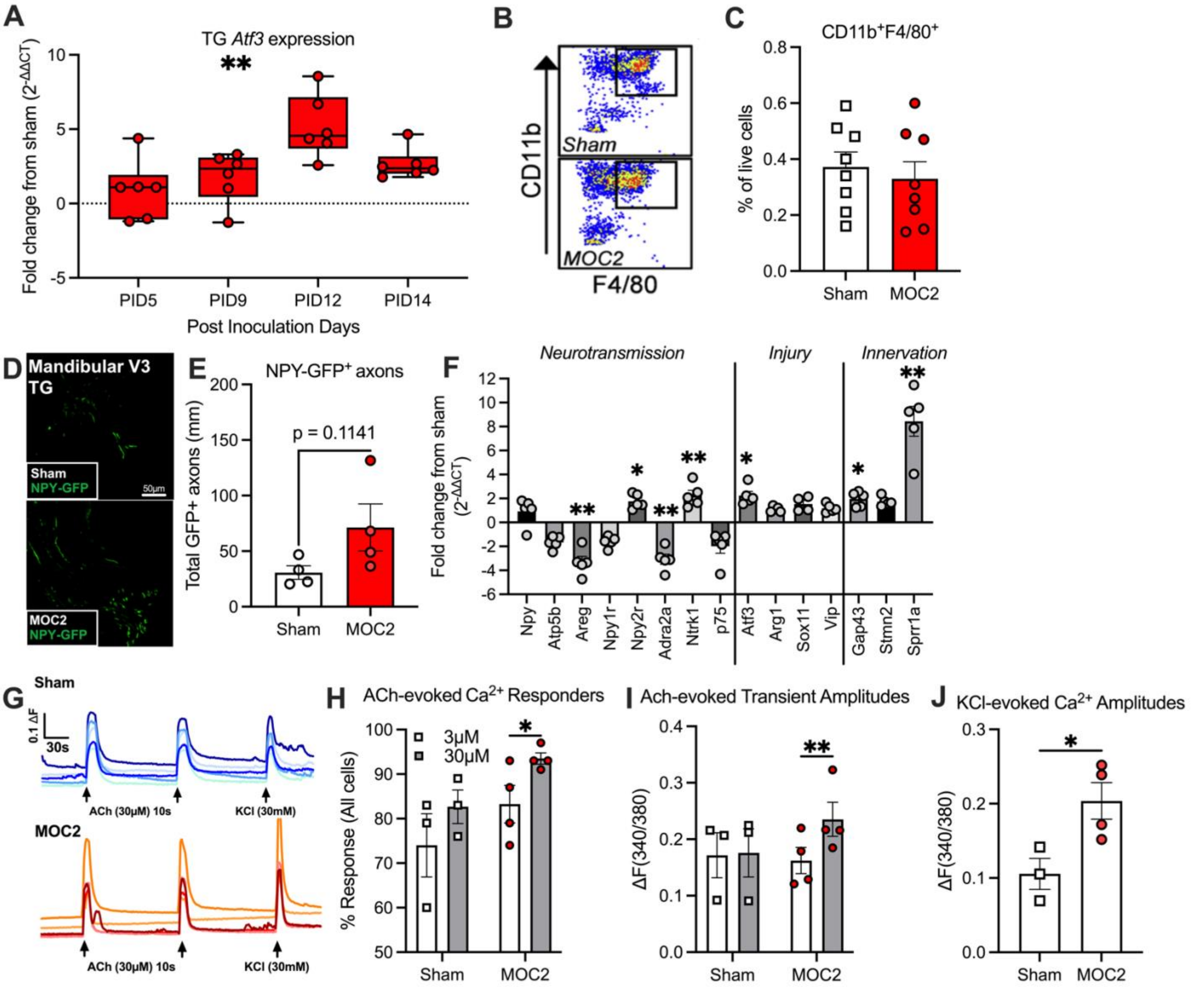


**Supplemental Figure 5: Sensory nerve injury in tumor bearing mice**

(A) Time-course qPCR analysis of *Atf3* expression in whole trigeminal ganglia (*93*) from MOC2 tumor-bearing mice. *Atf3* expression peaked at PID12, coinciding with the peak in spontaneous pain behavior. n=3/sex/group, Two-Way ANOVA, Cancer by time interaction, **$p<0.01$

(B-C) Flow cytometric analysis of dissociated trigeminal ganglia cells from sham and MOC2 mice at PID12. Representative scatter plots of $CD11b^{+}F4/80^{+}$ macrophages as a percent of live cells. n=4/sex/group, Independent T-test, $p>0.05$.

(D-E) Representative images of NPY-GFP+ sympathetic fibers in the V3 branch of the TG from Sham (Top) and MOC2 (Bottom) tumor-bearing mice. Pooled quantification of NPY-GFP+ fiber density reveals increased sympathetic fiber presence in MOC2 mice compared to sham controls. n=2/sex/group, Independent T-test, $p>0.05$.

(F) qPCR Gene Expression as a fold change from sham of neurotransmission, injury and sprouting associated genes in the SCG from sham and MOC2 tumor bearing mice. Analysis revealed upregulation of neurotransmission (*Areg, Npyr2, Adra2a, Ntrk1, p75*) and sprouting (*Gap43, Sprr1a*) associated genes. n=3/sex/group. Two-Way ANOVA, Cancer by gene interaction, *$p<0.05$, **$p<0.01$.

(G) Representative traces of reproducible acetylcholine (30uM)-evoked Ca2+ transients and KCl depolarization-evoked Ca2+ transient in dissociated SCG neurons from sham male mouse.

(H-I) Percent of acetylcholine (Ach, 3uM, 30uM)-responsive neurons and amplitude of evoked response in dissociated SCG from sham and MOC2 mice F) Amplitude of depolarization-evoked transient via 30mM KCl. SCG neurons from MOC2 mice are more sensitive to Ach and generate a larger $Ca^{2+}$ response to depolarization compared to SCG neurons from sham mice. n=1-2/sex/group. Two-way ANOVA; Cancer by drug interaction, *$p<0.05$, **$p<0.01$

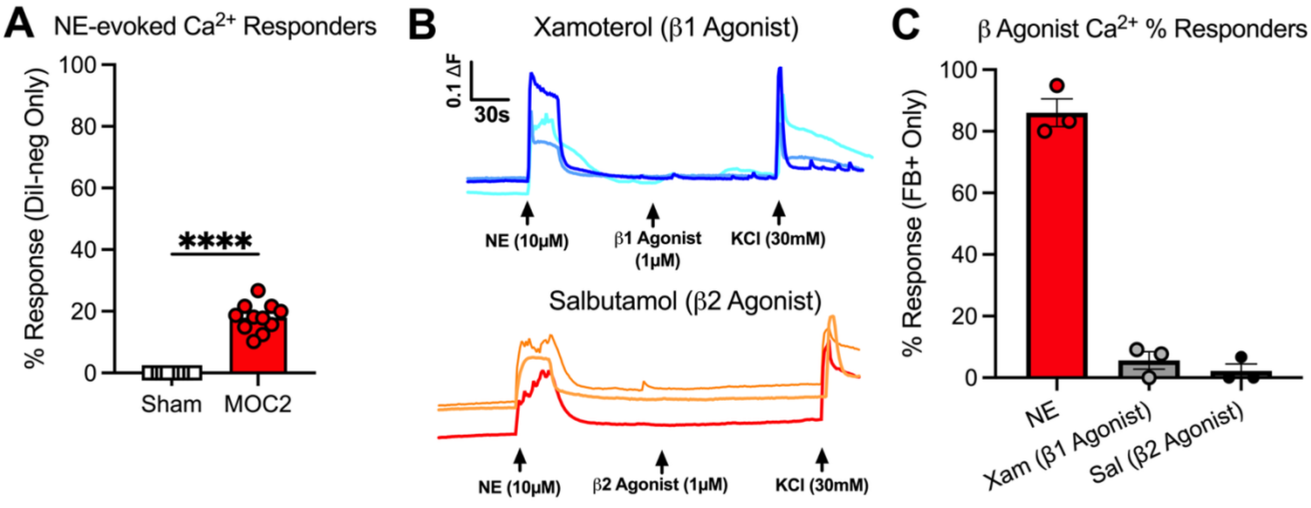


**Supplemental Figure 6: Adrenergic receptor pharmacology**
(A) Percent NE responders, tracer negative TGN’s, demonstrates about 20% responders. n=5-6/sex/group, Independent T-test, ****p<0.0001.

(B-C) Calcium imaging of tongue-innervating TGNs revealed using Beta1 and Beta 2 adrenergic pharmacology did not elicit $Ca^{2+}$ transients unlike NE. The percent responders to B1 and B2 agonist was below 5%. n=3/group, One-Way ANOVA, p>0.05.

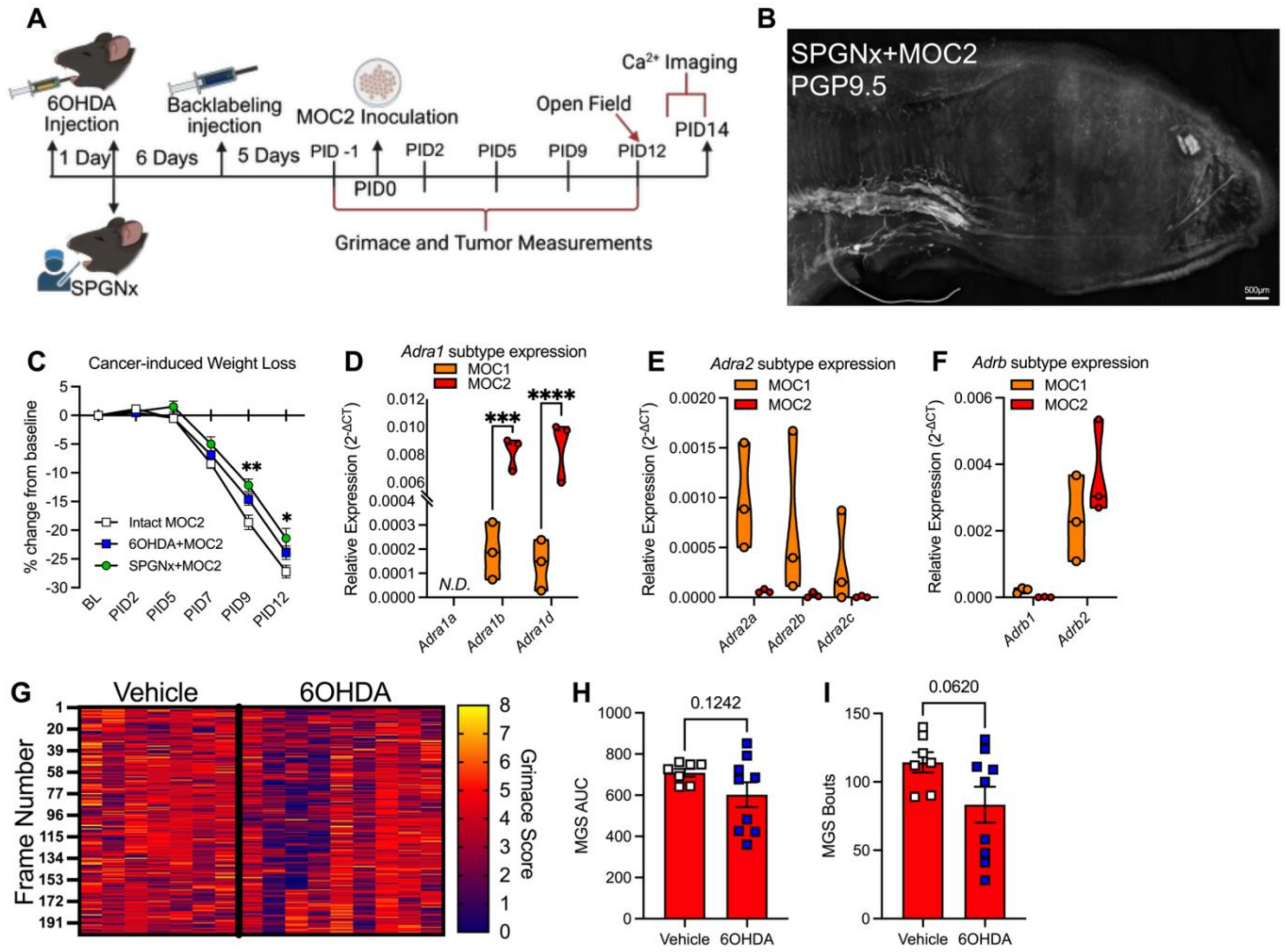


**Supplemental Figure 7: Sympathectomy decreased nociceptive behaviors**

(A) Timeline schematic of procedure for 6OHDA and SPGNx denervated mice experiments. Briefly, mice underwent surgical removal of bilateral superior cervical ganglia (SCG) and allowed to recover for 6 days prior to injection of retrograde tracer, DiI, into the anterior tongue (i.e. backlabeling injection). As an alternative to surgery, mice received two injections of 6OHDA (6 mg/kg) into the anterior (Day 1) and posterior (Day 2) to chemically denervate the tongue; again mice recovered for 6 days prior to backlabeling injection. Sympathectomized mice were inoculated with MOC2 ($2x10^4$) after 5 days. Starting on post inoculation day (PID) 1, weight changes, tumor growth and nociceptive behaviors (grimace) were tracked every 2-3 days; open field test (OFT) was executed on PID 12.

(B) Representative immunostaining of PGP9.5 (pan-neuronal marker) in a 300μm sagittal tongue section from a MOC2 tumor bearing mouse with surgical sympathectomy to demonstrate intact remaining sensory and non-SNS motor innervation

(C) Tongue tumor-induced weight loss in MOC2 mice with intact sympathetic innervation (pooled sham surgery and vehicle injection) compared to mice with chemical (6OHDA) and surgical sympathectomy (SPGNx); Mice with intact innervation lost more weight compared to SPGNx mice. n=3-5/sex/group, Two-way ANOVA, treatment by time interaction, *$p<0.05$, **$p<0.01$.

(D-F) quantitative real-time PCR gene expression as relative expression ($2^{-deltaCT}$) for each adrenergic receptor subtype in both MOC1 and MOC2 cell lines. Analysis revealed increased expression of *Adra1b* and *Adra1d* in MOC2 compared to MOC1, with *Adra2* and *Adrb* subtypes demonstrating no significant differences. n=3 cell line passages/group, Independent T-Test within gene, ***$p<0.005$, ****$p<0.0001$. N.D. = not detected.

(G-I) Grimace assay analysis in 6OHDA-treated sham and MOC2 mice. Heatmap (D), area under the curve grimace score (E), and grimacing bouts (F) at PID12 demonstrate a trend in less nociceptive behavior in MOC2 mice lacking sympathetic innervation in the tongue. n=3-5/sex/group, Independent T-Test, $p>0.05$.

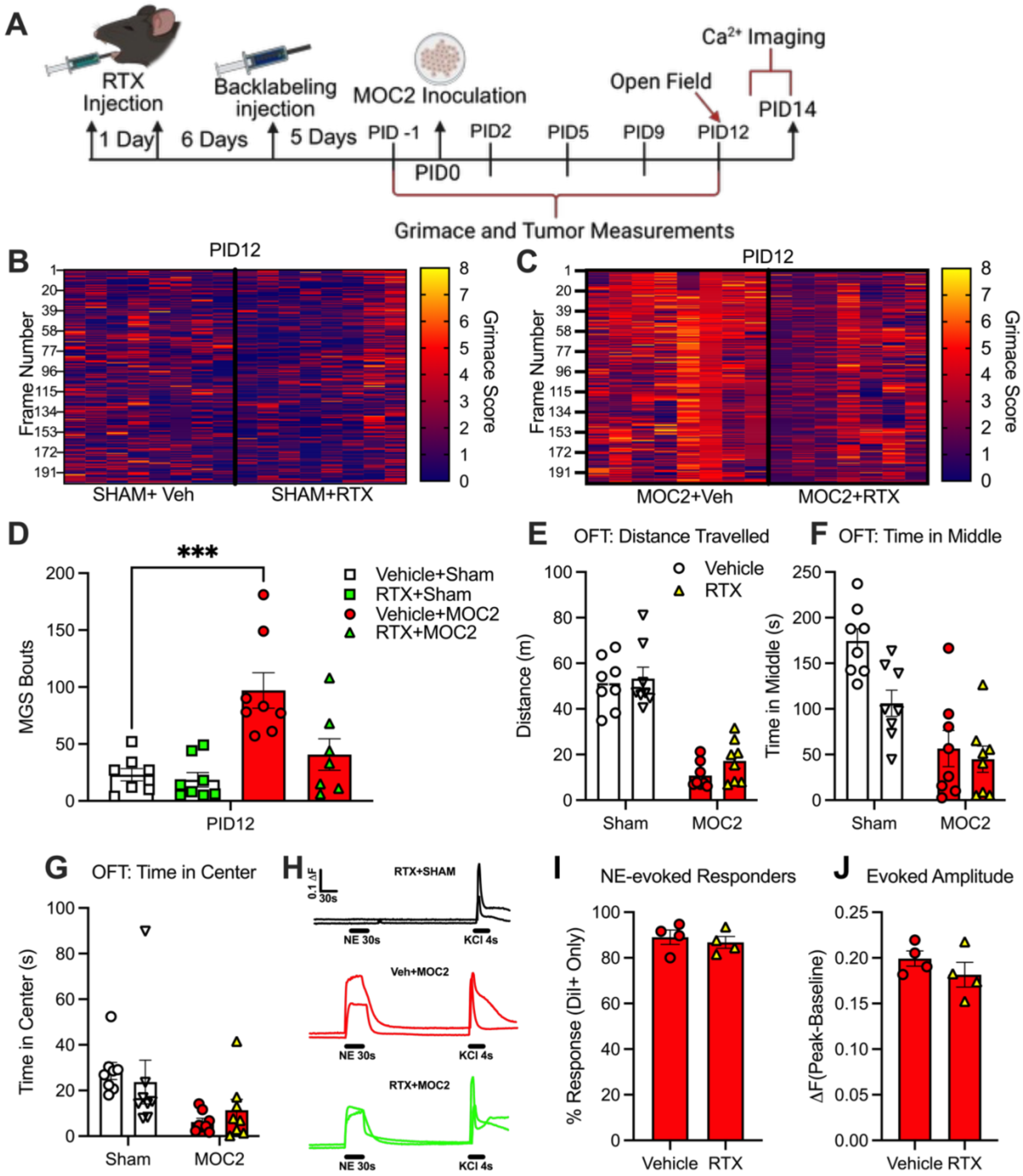


**Supplemental Figure 8: Sensory neuron ablation does not affect anxiety-like behaviors or acquired adrenergic sensitivity**

A) Timeline schematic of procedure for RTX-mediated denervation experiments. Briefly, mice received two injections of RTX (50 ng in 30 µl) into the anterior (Day 1) and posterior (Day 2) tongue to chemically denervate sensory tongue innervation. Mice were allowed to recover for 6 days prior to injection of retrograde tracer, DiI, into the anterior tongue (i.e. backlabeling injection). Denervated mice were inoculated with MOC2 ($2x10^4$) after 5 days. Starting on post inoculation day (PID) 1, weight changes, tumor growth and nociceptive behaviors (grimace) were tracked every 2-3 days; open field test (OFT) was executed on PID 12.

B) Heatmap visualization of grimace scores in sham mice treated with vehicle or resiniferatoxin (RTX) shows no change in spontaneous grimacing behavior following RTX administration.

(C-D) Heatmap visualization of grimace scores in MOC2 mice treated with vehicle or resiniferatoxin (RTX) at PID 12 shows a decrease in spontaneous grimacing behavior following RTX administration. Pooled quantification of grimacing frames (i.e. bouts) defined as ≥1.5 standard deviations from baseline n=3-4/sex/group, One-way ANOVA, *p<0.05.

(E-G) Open Field Test was used to evaluate anxiety-related behavior. Tumor-bearing mice exhibited anxiety-like phenotypes, including (D) reduced total distance traveled, (E) less time spent in the middle zone, and (F) less time spent in the center zone of the arena. These anxiety-related measures were not significantly altered by RTX treatment. n=4/sex/group, Two-way ANOVA, $p>0.05$.

(H-J) Calcium imaging of tongue-innervating TGNs revealed that RTX treatment did not induce adrenergic sensitivity in Sham mice. The percentage of NE-responsive neurons and the amplitude of calcium transients in response to 10 μM norepinephrine were not significantly different between RTX- and vehicle-treated groups in tumor bearing mice. n=2/sex/group, Independent T-test, $p>0.05$.

**Supplemental Table 1: Multivariable Regression: Pain Effects on Log of NE**

| Measure | Beta | 95% CI | p-value | q-value |
|---|---|---|---|---|
| BPI Mean Pain Severity | 0.16 | 0.059, 0.261 | 0.002 | 0.035 |
| BPI Mean Pain Interference | 0.174 | 0.068, 0.280 | 0.002 | 0.027 |
| OCPQ Composite Score | 0.003 | 0.002, 0.004 | <0.001 | <0.001 |
| OCPQ Function-evoked Pain | 0.005 | 0.002, 0.008 | 0.001 | 0.023 |
| OCPQ Spontaneous Pain | 0.007 | 0.004, 0.010 | <0.001 | <0.001 |

Note. Effects are after controlling for sex, T-stage, age, and PNI status

**Supplemental Table 2: Univariable Logistic Regression: Predictor Effects on PNI Status**

| Predictor | N | OR | 95% CI | p-value | q-value |
|---|---|---|---|---|---|
| PHQ-8 Sum | 90 | 1.02 | 0.94, 1.12 | 0.6 | >0.9 |
| GAD-7 Sum | 90 | 0.99 | 0.89, 1.09 | 0.8 | >0.9 |
| Norepinephrine (NE) | 58 | 1 | 1.00, 1.01 | 0.034 | 0.03 |
| Sex | 90 | | | 0.4 | >0.9 |
| F | | — | — | | |
| M | | 1.54 | 0.62, 4.08 | | |
| Age | 90 | 1 | 0.96, 1.03 | 0.8 | >0.9 |
| Primary Tumor-Stage | 90 | | | 0.3 | >0.9 |
| T1/T2 | | — | — | | |
| T3/T4 | | 1.65 | 0.67, 4.08 | | |
| Nodal Status | 90 | | | 0.003 | 0.043 |
| N0/N1 | | — | — | | |
| N2/N3 | | 4.24 | 1.61, 11.6 | | |
| Extra Capsular Spread | 88 | | | 0.071 | 0.6 |
| No | | — | — | | |
| Yes | | 2.58 | 0.92, 7.29 | | |

**Supplemental Table 3: Multivariable Logistic Regression: Pain Effects on PNI Status**

| Measure | N | OR | 95% CI | p-value |
|---|---|---|---|---|
| BPI Mean Pain Severity | 58 | 1.22 | 0.92, 1.65 | 0.2 |
| BPI Mean Pain Interference | 57 | 1.06 | 0.78, 1.43 | 0.7 |
| OCPQ Composite Score | 58 | 1 | 1.00, 1.01 | 0.027 |
| OCPQ Function-evoked Pain | 58 | 1.01 | 1.00, 1.02 | 0.12 |
| OCPQ Spontaneous Pain | 58 | 1.01 | 1.00, 1.02 | 0.011 |

*Note*. Effects are after controlling for NE, N-stage, T-stage, and sex

**Supplemental Table 4: Taqman Assays for single cell and whole ganglia quantitative PCR**

| Gene Name | Protein name (target) | Thermo Fisher Scientific Taqman Assay |
|---|---|---|
| Adra1a | α1A-adrenergic receptor | Adra1a_Mm00442668_m1 |
| *Adra1b* | α1B-adrenergic receptor | Adra1b_Mm00431685_m1 |
| *Adra1d* | α1D-adrenergic receptor | Adra1d_Mm01328600_m1 |
| *Adra2a* | α2A-adrenergic receptor | Adra2a_Mm07295458_s1 |
| *Adra2b* | α2B-adrenergic receptor | Adra2b_Mm00477390_s1 |
| *Adra2c* | α2C-adrenergic receptor | Adra2c_Mm00431686_s1 |
| *Adrb1* | β1-adrenergic receptor | Adrb1_Mm00431701_s1 |
| *Adrb2* | β2-adrenergic receptor | Adrb2_ Mm02524224_s1 |
| *Areg* | Amphiregulin | Areg_Mm01354339_m1 |
| *Arg1* | Arginase-1 | Arg1_Mm00475988_m1 |
| *Atf3* | Activating transcription factor 3 | Atf3_Mm00476032_m1 |
| *Atp5b* | ATP synthase subunit beta | Atp5b_Mm01160389_g1 |
| *Gap43* | Growth associated protein 43 | Gap43_Mm00500404_m1 |
| *Npy* | Neuropeptide Y | Npy_Mm01410146_m1 |
| *Npy1r* | Neuropeptide Y receptor type 1 | Npy1r_Mm00650798_g1 |
| *Npy2r* | Neuropeptide Y receptor type 2 | Npy1r_Mm00650798_g1 |
| *Ntrk1* | Tropomyosin receptor kinase A (TrkA) | Ntrk1_Mm01219406_m1 |
| *p75* | p75 neurotrophin receptor (NGFR) | Ngfr_Mm00446296_m1 |
| *Sox11* | SRY-box transcription factor 11 | Sox11_Mm01281943_s1 |
| *Sprr1a* | Small proline-rich protein 1A | Sprr1a_Mm01962902_s1 |
| *Stmn2* | Stathmin-2 (SCG10) | Stmn2_Mm00600432_m1 |
| *Vip* | Vasoactive intestinal peptide | Vip_Mm00660234_m1 |
